# The Origin of Cysteine and its Catabolism in Mammalian Tissues and Tumors

**DOI:** 10.1101/2022.08.26.505162

**Authors:** Sang Jun Yoon, Joseph A. Combs, Aimee Falzone, Nicolas Prieto-Farigua, Samantha Caldwell, Hayley D. Ackerman, Elsa R. Flores, Gina M. DeNicola

## Abstract

Cysteine plays critical roles in cellular biosynthesis, enzyme catalysis, and is an essential contributor to redox metabolism. While cultured cells are highly dependent on exogenous cystine for proliferation and survival, how diverse tissues obtain and use cysteine in vivo has not been characterized. We comprehensively interrogated cysteine metabolism in normal murine tissues and the cancer that arise from them using stable isotope ^13^C-serine and ^13^C-cystine tracing. We found that de novo cysteine synthesis was highest in normal liver and pancreas and absent in lung tissue. In tumors, cysteine synthesis was either inactive or downregulated during tumorigenesis. By contrast, cystine uptake and metabolism to downstream metabolites was a universal feature of normal tissues and tumors. Differences in cysteine catabolism were evident across tumor types, including glutathione synthesis. Thus, cystine is a major contributor to the cysteine pool in tumors and cysteine catabolic pathways are differentially active across tumor types.

## INTRODUCTION

The non-essential, thiol-containing amino acid cysteine is an essential source of sulfur for the synthesis of diverse cellular factors that play important biological functions in maintaining redox homeostasis, enzyme catalysis, and electron transfer (Ward and DeNicola, 2019). Cysteine is partitioned into various downstream metabolic pathways including glutathione biosynthesis and cysteine oxidation. The tripeptide antioxidant glutathione is the most abundant intracellular antioxidant (Winterbourn and Hampton, 2008) and is synthesized from cysteine, glutamate, and glycine in two steps mediated by glutamate-cysteine ligase (GCL) and glutathione synthetase (GSS) (Anderson, 1998). Glutathione synthesis is regulated by the rate limiting enzyme GCL (Lu, 1999), which consists of a catalytic subunit (GCLC) and a modifier subunit (GCLM), which relieves the feedback inhibition of GCLC by glutathione. Glutathione synthesis is induced by oxidative stress, which stabilizes nuclear factor erythroid 2–related factor 2 (NRF2) to induce the transcription of both GCLC and GCLM and a battery of antioxidant enzymes that detoxify reactive oxygen species (ROS) (Harris and DeNicola, 2020).

Tumorigenesis is accompanied by an increased demand for cysteine to deal oxidative stress (Pavlova and Thompson, 2016; Pavlova et al., 2022). Many cancers upregulate the system x_c_^-^ cystine/glutamate antiporter (xCT) to maintain the intracellular cysteine pool and promote entry of cysteine into glutathione synthesis (Zhang et al., 2022). This is achieved by mutations in oncogenes/tumor suppressors that regulate xCT expression, including Kelch-like ECH-associated protein 1 (KEAP1)/NRF2 (Kang et al., 2019; Sasaki et al., 2002) and p53 (Jiang et al., 2015), or regulate xCT activity (Gu et al., 2017; Lien et al., 2017; Mukhopadhyay et al., 2021; Tsuchihashi et al., 2016). Moreover, xCT expression is induced by amino acid starvation via activating transcription factor 4 (ATF4) (Sato et al., 2004), thereby ensuring adequate cysteine availability in nutrient poor conditions. There are many lines of evidence that extracellular cystine is the primary supply of the intracellular cysteine pool to support the cellular redox state. Insufficient cystine availability induces iron-dependent lipid peroxidation, leading to a form of cell death known as ferroptosis (Dixon et al., 2012; Kang et al., 2021; Zhang et al., 2020). Pharmacological inhibition of xCT induces ferroptosis of cancer cells (Dixon *et al*., 2012; Zhang et al., 2019), and enzyme-based cystine degradation has shown efficacy against several in vivo cancer models (Badgley et al., 2020; Cramer et al., 2017; Zhang *et al*., 2019).

Beyond cystine uptake, the intracellular cysteine pool can be sustained by the transsulfuration pathway in the liver, although its contribution to other tissues is less clear (Combs and DeNicola, 2019). Transsulfuration is catalyzed by cystathionine β-synthase (CBS) and cystathionine γ-lyase (CSE) and mediates both irreversible homocysteine removal and de novo cysteine synthesis, with serine donating the carbon backbone and homocysteine donating the sulfur to cysteine (Stipanuk, 2004b). While both CBS and CSE are broadly expressed (Mudd et al., 1965), the contribution of transsulfuration to the cysteine pool in tumors is poorly characterized. Prior studies have found a small contribution of this pathway to the cysteine pool, which could protect against cystine starvation in Ewing’s Sarcoma and neuroblastoma cell lines (Zhang et al., 2021; Zhu et al., 2019). By contrast, we found no contribution of the transsulfuration pathway to the cysteine and glutathione pool in non-small cell lung cancer (NSCLC) cell lines, which robustly die by ferroptosis under cystine starvation (Kang *et al*., 2021). Moreover, the contribution of the transsulfuration pathway to the cysteine pools of non-hepatic tissues and tumors in vivo has not been investigated.

In this study, we comprehensively investigated the contribution of the both the transsulfuration pathway and exogenous cyst(e)ine to the cysteine pool and downstream metabolites in nine different healthy mouse tissues and tumors of the lung, pancreas and liver. We found limited contribution of transsulfuration to the cysteine pool of non-hepatic tissues, in contrast to robust contribution from exogenous cyst(e)ine. Moreover, tumors from transsulfuration capable tissues downregulated this pathway, while tissues lacking transsulfuration activity generated transsulfuration deficient tumors. Finally, we characterized cysteine catabolism to glutathione and taurine across tissues and tumors, which demonstrated complex patterns associated with enzyme expression and substrate availability.

## RESULTS

### Cultured Cancer Cell Lines Lack of De Novo Cysteine Synthesis Capacity

In our previous study, we found that NSCLC cysteine pools are not supported by transsulfuration, resulting in cumulative cell death following extracellular cystine starvation (Kang *et al*., 2021). To evaluate the origin of cysteine more broadly in cancer cells in culture, we examined de novo cysteine synthesis from ^13^C_3_-serine to cysteine through the transsulfuration pathway in a panel of small cell lung cancer (SCLC), pancreatic ductal adenocarcinoma (PDAC) and hepatocellular carcinoma (HCC) cell lines. Incubation with ^13^C_3_-serine as an extracellular serine source for 4 hours resulted in almost complete labeling of the intracellular serine pool (Figure 1A), and metabolism to cystathionine (Figure 1B). We found that SCLC and NSCLC showed higher cystathionine labeling in this time period than HCC and PDAC cell lines (Figure 1B). However, regardless of cancer type, labeling of cysteine from serine was absent in all cell lines, suggesting a bottleneck at cysteine synthesis from cystathionine (Figure 1C). To examine whether impaired cysteine synthesis is a consequence of lack of transsulfuration enzyme expression, we performed immunoblotting for CBS and CSE, which mediate the first and second steps of transsulfuration (Figure 1D), respectively. Interestingly, while some PDAC cell lines lacked CBS expression and some HCC cell lines lacked CSE expression, almost all cancer cell lines investigated expressed both enzymes despite being unable to synthesize cysteine (Figure 1D).

**Figure 1.**
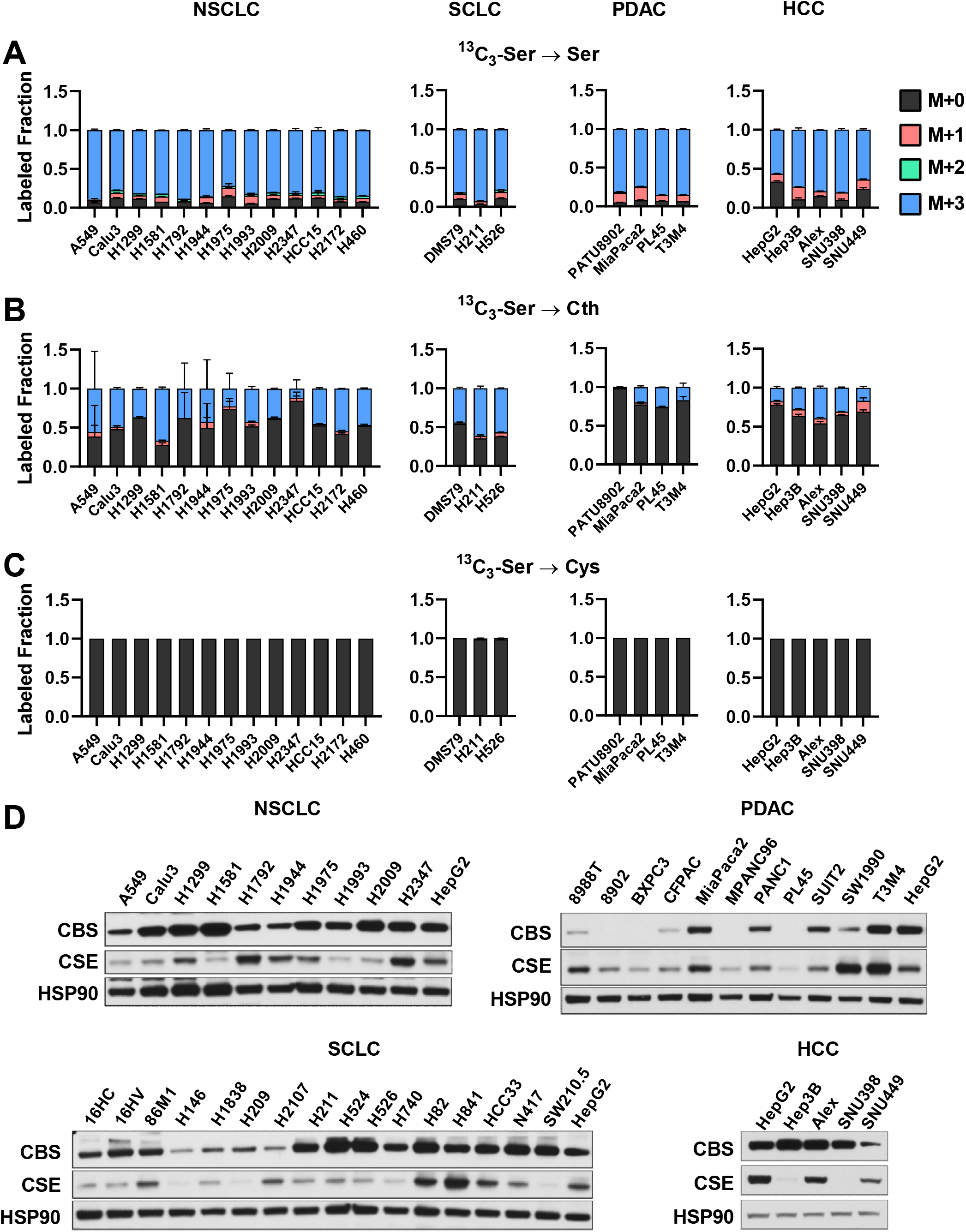
Cultured Cancer Cell Lines Lack of De Novo Cysteine Synthesis Capacity. (A-C) Analysis of de novo cysteine synthesis in cultured NSCLC, SCLC, PDAC, and HCC cell lines with ^13^C_3_-serine tracing. Cell lines were incubated with ^13^C_3_-serine containing media for 4 hours, followed by analysis of the fraction labeling in (A) serine, (B) cystathionine and (C) cysteine. Data are presented as mean ± SD and N=3 biological replicates for each cell line. (D) Immunoblotting for the transsulfuration enzymes cystathionine β-synthase (CBS) and cystathionine γ-lyase (CSE). HSP90 was used for the loading control and HepG2 was used for relative comparison between different membranes. Ser, serine; Cth, cystathionine; Cys, cysteine; Gly, glycine; GSH, glutathione; γ-Glu-Cys, γ-glutamylcysteine.

Prior studies have suggested that generation of homocysteine via the methionine cycle is a critical limiting factor for transsulfuration flux (Ye et al., 2017; Zhu *et al*., 2019). To examine whether transsulfuration may be substrate-limited, we examined whether excess cystathionine or homocysteine could rescue viability under cystine starvation in a panel of NSCLC cell lines (Figure S1A). We observed a full rescue of viability by either substrate, which was reversed by treatment with propargylglycine, an irreversible inhibitor of CSE, demonstrating the requirement for transsulfuration for this rescue (Figure S1B). To confirm that these substrates were actively contributing to the cysteine and glutathione pools, we cultured cells with ^13^C_3_-serine for 24 hours under cystine replete or starved conditions in the presence or absence of cystathionine or homocysteine. Homocysteine treatment elevated both homocysteine and cystathionine levels within cells, with cystathionine demonstrating M+3 labeling from serine (Figure S1C). Because cystathionine already contains serine carbons, ^13^C_3_-serine was not useful for assaying transsulfuration under cystathionine treatment conditions. Nevertheless, we observed that cystathionine treatment was more effective at elevating cystathionine and did not alter homocysteine levels as expected. Both treatments elevated cysteine levels under cysteine starved conditions, although levels were still much lower than replete conditions. Despite this, glutathione levels were completely restored, with M+2 labeling (via glycine) demonstrating active de novo synthesis despite the lack of exogenous cystine. Moreover, M+3 labeling of cystathionine, cysteine, and glutathione was evident in homocysteine treated conditions, demonstrating active cysteine synthesis from serine. These results indicate that under these culture conditions transsulfuration enzymes are substrate limited for the synthesis of cysteine.

### Contribution of De Novo Cysteine Synthesis to the Cysteine Pool Varies across Healthy Mouse Tissues

Transsulfuration activity is known to be high in healthy liver (Mudd *et al*., 1965), but the contribution of this pathway to the cysteine pool across diverse tissues is not known. To this end, we infused healthy C57BL/6J mice with 1-[^13^C_1_]-serine for 4 hours via the jugular vein to label intracellular intermediates in the transsulfuration and glutathione synthesis pathways (Figure 2A). We analyzed their fraction labeling (Figure 2B-G) and total levels (Figure S2A-S2F) in nine different tissues (liver, pancreas, kidney, heart, thymus, spleen, lung, cerebellum, and brain) and serum by mass spectrometry. The resulting total signal intensity of intermediates revealed that liver and pancreas are the most cystathionine abundant tissues, while pancreas and kidney have a larger cysteine pool than the others (Figure S2C and S2E). Four hours of 1-[^13^C_1_]-serine infusion labeled around 50% of circulating serine (Figure 2B) and this time frame was sufficient to detect ^13^C label in cysteine and glutathione (Figure 2E, F). Within each tissue, fraction labeling was normalized to labeling in the serine pool to evaluate the fractional contribution of serine to downstream intermediates. Like what we observed in cultured cell lines (Figure 1B), cystathionine labeling was detected in all tissues (Figure 2D), demonstrating that CBS was active. When normalized to labeling of intracellular serine, many tissues demonstrated very high cystathionine labeling including heart (98%), liver (84%), lung (76%), and spleen (65%) (Figure S2G). Moreover, brain had a fraction labeling >100%, suggesting uptake of cystathionine from the circulation. Analysis of cysteine labeling across tissues revealed robust de novo cysteine synthesis in liver tissue (25% labeled), and the second highest labeling was observed in pancreas (3%), with other tissues deriving less than 3% of their cysteine from serine (Figure 2H). Given the high fractional contribution of serine to cystathionine across tissues (Figure S2G), these results suggest that the cleavage of the bond between sulfur and the gamma carbon by CSE is a bottle neck of de novo cysteine synthesis.

**Figure 2.**
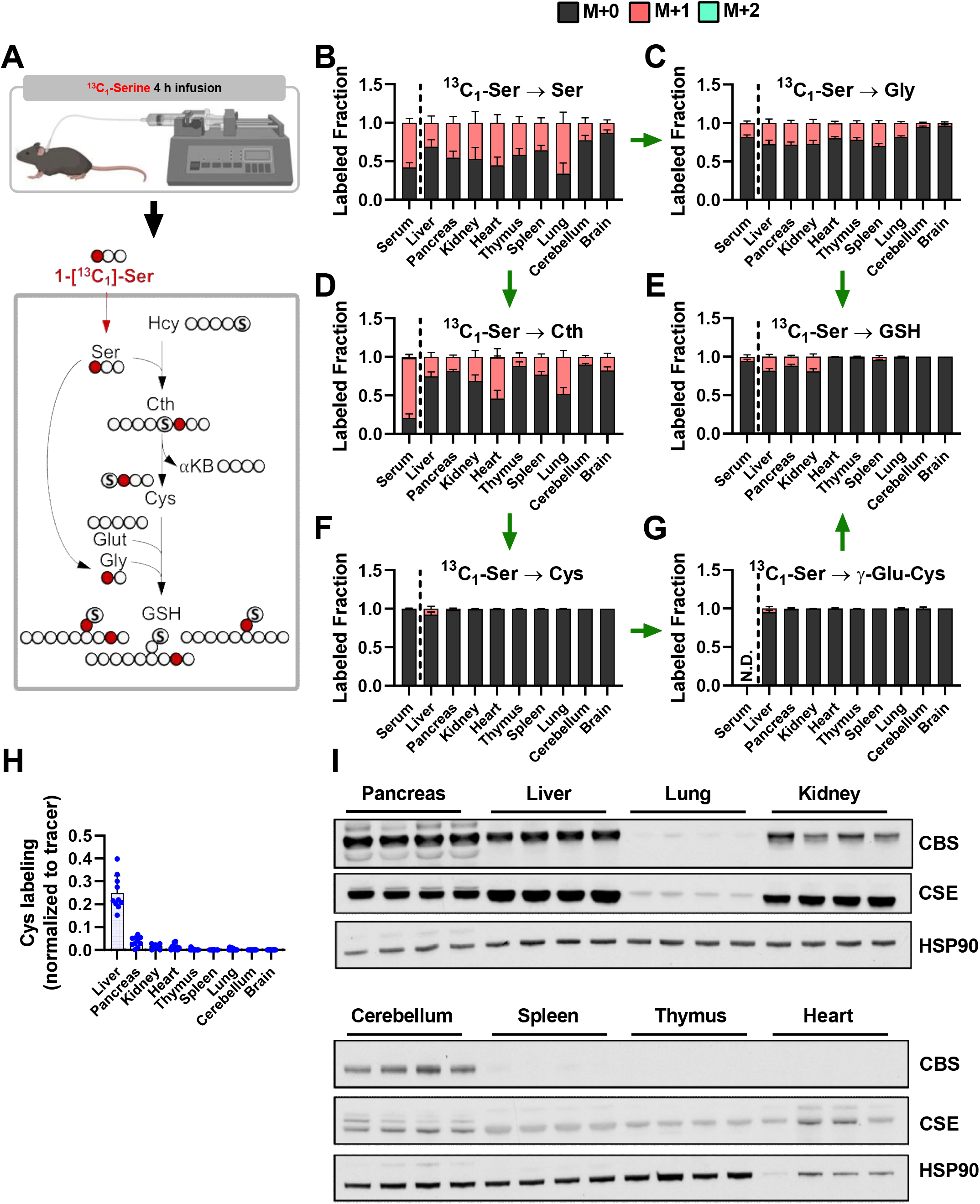
Contribution of De novo Cysteine Synthesis to the Cysteine Pool Varies across Healthy Mouse Tissues. (A) Schematic depicting 1-[^13^C_3_]-serine infusion and its metabolism via the transsulfuration pathway and glutathione synthesis pathways. (B-G) Healthy C57BL6J mice were infused with 1-[^13^C_3_]-serine, followed by analysis of the fraction labeling in (B) serine, (C) glycine, (D) cystathionine, (E) glutathione, (F) cysteine and (G) γ-glutamylcysteine. For (B-G), data are presented as mean ± SD and N=10 mice (5 male, 5 female). (H) Fractional contribution of serine to intracellular cysteine synthesis in each tissue from B-G. Cysteine labeling was normalized to the fraction labeling of serine in each tissue. (I) Immunoblots of cystathionine β-synthase (CBS) and cystathionine γ-lyase (CSE) for each tissue. HSP90 was used for the loading control.

Next, we examined CBS and CSE expression levels across tissues to examine their association with cysteine synthesis (Figure 2I). We found that liver and pancreas both have high expression of CBS, with the liver demonstrating higher CSE expression than pancreas (Figure 2I). Even when accounting for cystathionine labeling, liver had much higher synthesis of cysteine compared to pancreas, suggesting that second step of transsulfuration mediated by CSE was more active (Figure 2H and S2G), and the expression of CSE and additional factors may be key regulators of cysteine synthesis (Figure 2I). Tissues lacking cysteine labeling generally had low expression of both CBS and CSE, including lung, cerebellum, thymus and spleen (Figure 2F, 2H and 2I). Kidney had high expression of CSE, but lower CBS expression than liver and pancreas (Figure 2I), which likely explained its lower cysteine labeling (1.5%, Figure 2H). Collectively, these results demonstrate that transsulfuration of serine to cysteine is a minor contributor to the cysteine pool in most non-hepatic tissues and the bottleneck is the CSE step.

1-[^13^C_1_]-serine infusion for 4 hours was also sufficient to evaluate the glutathione synthesis pathway downstream of cysteine metabolism. Evaluation of total metabolite pools revealed that liver had the highest total signal of glutathione (Figure S2D), consistent with its established role in glutathione synthesis to supply the circulating pool (Ookhtens and Kaplowitz, 1998). Liver, pancreas, and kidney and spleen demonstrated detectable labeling from serine (Figure 2E). When normalized to labeling of intracellular serine, liver demonstrated the highest labeling (60%), followed by kidney (43%), pancreas (26%), and spleen (12%) (Figure S2H). However, because 1-[^13^C_1_]-serine can label glutathione via both cysteine and glycine, with both resulting in M+1 labeling, this labeling likely overestimates the contribution of cysteine (Figure 2F). To directly examine the entry of serine-derived cysteine into the glutathione synthesis pathway, we examined labeling in γ-glutamylcysteine (Figure 2G). In agreement with cysteine labeling, only liver showed a substantial fraction of γ-glutamylcysteine labeling (5%). Additionally, M+2 labeling of glutathione was absent across all tissues (Figure 2E), further supporting that serine-derived cysteine comprises a minor proportion of glutathione.

### Cyst(e)ine Supplies the Cysteine Pool in All Tissues

To directly assay cysteine metabolism to downstream metabolites, we infused healthy C57BL6/J mice with ^13^C_6_-cystine for 4 hours (Figure 3A). Serum and tissues were collected and analyzed by LC-MS based metabolomics to examine the total signal (Figure S3A-S3F) and labeled fraction (Figure 3B-3G) of intermediates within the glutathione synthesis and cysteine oxidation pathways. Interestingly, despite infusion with pure ^13^C_6_-cystine, cystine formed mixed disulfides within the serum and tissues to form a substantial fraction of M+3 cystine (Figure 3B). Importantly, M+3 cysteine was also detected in the serum, precluding our ability to determine uptake as cystine vs. cysteine (Figure 3D and S3C). Cysteine synthesis low tissues including heart, thymus, and lung demonstrated the highest fraction labeling of cysteine from ^13^C_6_-cystine, while the lowest fraction labeling was observed in cysteine synthesis high pancreas and kidney (Figure 3D). Immunoblotting for the expression of the cystine/glutamate antiporter revealed highest expression in pancreas (Figure 3J), which also had the highest total cysteine levels (Figure S3C). However, xCT expression largely did not correlate with cysteine levels or labeling across tissues, suggesting other transporters for cystine or cysteine may mediate import, or additional mechanisms of xCT regulation may play a role. Indeed, we examined glutamate levels across tissues and found that brain and cerebellum levels were expectedly high (Figure S3G), consistent with the neurotransmitter function of this metabolite, which likely inhibits the ability of the xCT in cerebellum to import cystine (Figures 3B), which is very low in this tissue (Figure S3A). Moreover, these results support our finding that liver, pancreas, and kidney contribute to the cysteine pool through de novo cysteine synthesis (Figure 2H and Figure 3D).

**Figure 3.**
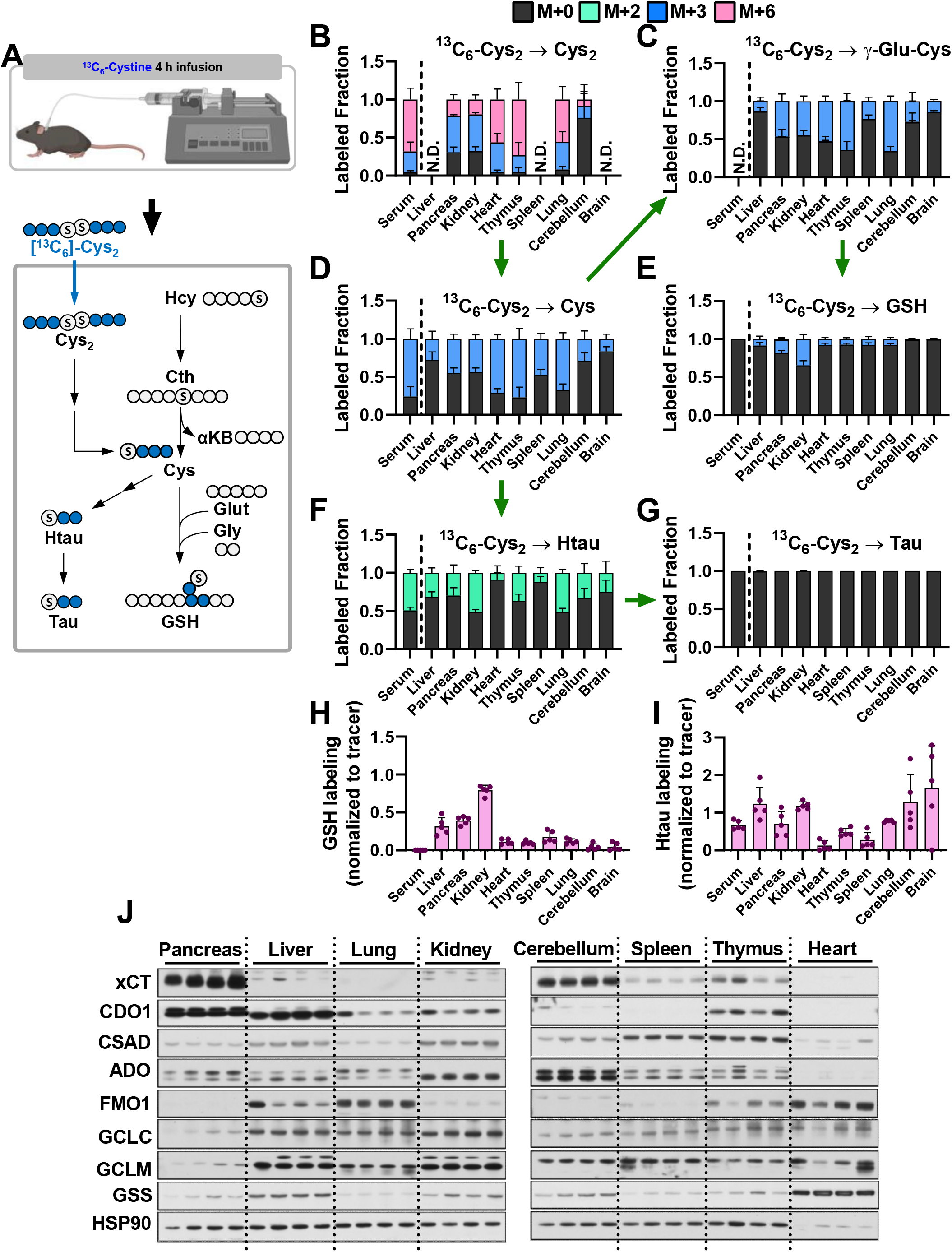
Cyst(e)ine supplies the cysteine pool in all tissues. (A) Schematic depicting ^13^C_6_-cystine infusion and its metabolism to glutathione and taurine. (B-G) Healthy C57BL6J mice were infused with ^13^C_6_-cystine, followed by analysis of the fraction labeling in (B) cystine, (C) γ-glutamylcysteine, (D) cysteine, (E) glutathione, (F) hypotaurine and (G) taurine. For (B-G), data are presented as mean ± SD and N=5 mice. (H-I) Fractional contribution of cystine to (H) glutathione and (I) hypotaurine in each tissue from B-G. Metabolite labeling was normalized to the fraction labeling of cysteine in each tissue. (J) Immunoblots of cystine/glutamate antiporter (xCT), cysteine dioxygenase type 1 (CDO1), cysteine sulfinate decarboxylase (CSAD), 2-aminoethanethiol (cysteamine) dioxygenase (ADO), glutamate-cysteine ligase catalytic subunit (GCLC), glutamate-cysteine ligase modifier subunit (GCLM), and glutathione synthetase (GSS) for each tissue. HSP90 is used for the loading control. Ser, serine; Cth, cystathionine; Cys, cysteine; Gly, glycine; GSH, glutathione; γ-Glu-Cys, γ-glutamylcysteine

Because all tissues demonstrated substantial labeling in the cysteine pool from ^13^C_6_-cystine, we were able to examine differential metabolism of cysteine to downstream metabolites across tissues. We found that kidney, pancreas, and liver demonstrate significantly higher glutathione labeling compared with the other tissues, with 79% labeling observed in kidney (Figure 3H). By contrast, brain and cerebellum demonstrated the lowest glutathione labeling at approximately 4%, suggesting glutathione synthesis may be very slow in these tissues (Figure 3E, 3H, and S3D). The rate-limiting step of glutathione synthesis depends on regulation of GCL activity, which is positively regulated cysteine availability and negatively regulated by glutathione (Lu, 1999). High glutathione labeling in liver and kidney was correlated with strong expression of both subunits of GCL (GCLC and GCLM) (Figure 3J), with the lower level of cysteine and higher level of glutathione potentially contributing to the higher labeling in the kidney (Figure 3H, S3C and S3D). While pancreas had low expression of GCLC and GCLM, its high level of xCT expression and cysteine levels and low levels of glutathione may contribute to high glutathione labeling (Figure 3H, 3J, S3C and S3D). For lung, cerebellum, spleen, and thymus, lower expression of GCLC and GCLM combined with relatively low cysteine and high glutathione likely contribute to lower levels of de novo glutathione synthesis (Figure 3H, 3J, S3C and S3D). Interestingly, the heart has higher expression of GSS compared to other tissues (Figure 3J), but has a low rate of glutathione synthesis (Figure 3H), raising the possibility that GSS has other functions. These results demonstrate diverse metabolism of cysteine to glutathione across murine tissues.

### Hypotaurine and Taurine Are Maintained by Crosstalk between Cysteine Catabolism and Transport

Next, we examined cysteine metabolism to taurine to determine if there are differences in cysteine entry into downstream pathways across tissues. Cysteine is metabolized to hypotaurine via the cysteine sulfinic acid pathway, mediated by cysteine dioxygenase type 1 (CDO1), or the cysteamine pathway downstream of Coenzyme A (CoA) breakdown, followed by oxidation to taurine. 4 hours of infusion with ^13^C_6_-cystine was sufficient to label hypotaurine in the serum and all tissues but was not sufficient to label taurine (Figure 3F and 3G). Interestingly, despite having low cysteine labeling and very low glutathione labeling, the brain and cerebellum had substantial hypotaurine labeling (Figure 3F). Consistent with prior reports (Stipanuk, 2004a; b), kidney and liver expressed CDO1 and cysteine sulfinic acid decarboxylase (CSAD) (Figure 3J). In addition to these tissues, we also observe strong CDO1 expression in pancreas, while cerebellum and thymus had low expression. CDO1 expression was undetectable in spleen and heart, which matched the lowest hypotaurine labeling in these two organs (Figure 3I, 3J). CSAD and 2-aminoethanethiol (cysteamine) dioxygenase (ADO) expression were more uniform across the tissues, except for heart, which lacked expression of both enzymes, and cerebellum, which had high ADO expression (Figure 3J). Interestingly, normalization of the hypotaurine labeled fraction to cysteine labeling revealed that liver, kidney, cerebellum, and brain demonstrated hypotaurine labeling higher than 100% (Figure 3I), suggesting that hypotaurine from the circulation may be contributing to the hypotaurine pool in these tissues. These results demonstrate that cysteine is prioritized for taurine synthesis in a subset of tissues, and the taurine pool may be supported by transport of taurine synthesis intermediates.

While the lack of labeling in taurine from ^13^C_6_-cystine precluded our ability to look directly at synthesis, there were interesting differences in hypotaurine and taurine levels across tissues that prompted us to look at the final step in taurine synthesis. Hypotaurine is enzymatically oxidized to generate taurine, although the enzyme responsible this reaction is not well defined. NAD-dependent hypotaurine dehydrogenase has been suggested as the responsible enzyme, but direct evidence is lacking (Sumizu, 1962). Recently, flavin-containing monooxygenase 1 (FMO1) was shown to mediate taurine biosynthesis from hypotaurine in vivo (Veeravalli et al., 2020). Immunoblotting revealed that liver, heart and lung had the highest FMO1 expression, while expression in pancreas was undetectable (Figure 3J). These patterns match taurine levels across tissues, with levels highest in liver and heart, with the levels in the pancreas almost an order of magnitude lower than heart (Figure S3F). These findings suggest FMO1 expressing tissues may have a greater capacity to metabolize hypotaurine to taurine.

### Tumorigenesis of Liver and Pancreas Induces Downregulation of De Novo Cysteine Synthesis

Given that cysteine is a crucial biomolecule which contains a sulfur moiety that facilitates redox homeostasis and energy transfer, cancers have been proposed to maintain their cysteine pool by rewiring de novo synthesis or cystine uptake (Zhang *et al*., 2022). To interrogate the source of cysteine in tumors in vivo, we first selected two genetically engineered mouse (GEM) models of liver and pancreatic cancer, since these two tissues demonstrated cysteine synthesis capacity, and examined whether cysteine synthesis capacity is maintained in tumors. A hydrodynamic tail vein injection model was used to generate *Myc; Trp53^-/-^* liver tumors for a HCC model and *LSL-Kras^G12D/+^; Trp53^flox/+^; p48-Cre* (KPC) mice were used for a PDAC model (Figure 4A). 1-[^13^C_1_]-serine tracing was performed as described for healthy mice. HCC demonstrated similar labeling in the serine pool compared to normal liver, and similar labeling in downstream metabolites cystathionine and glycine (Figure 4B). Labeling in the cysteine, γ-glutamylcysteine, and glutathione pools were lower in HCC compared to normal liver, although these differences were not significant (Figure 4B). PDAC demonstrated higher labeling in the serine pool, with similar labeling in downstream glycine and cystathionine pools (Figure 4C). However, labeling in both cysteine and γ-glutamylcysteine was absent. Although labeling was detected in glutathione, this was likely coming from glycine due to the absence of cysteine labeling (Figure 4C). When normalized to serine labeling within each tissue, we found that de novo cysteine synthesis decreased in both HCC and PDAC tumors compared with each control healthy tissue (Figure 4D). For HCC, the distribution appeared binary, with tumors either maintaining the labeling fraction of the parental tissue or having a substantially reduced fraction. For PDAC, cysteine labeling was completely absent (Figure 4D). Immunoblotting for CBS and CSE revealed a slight reduction in these proteins in HCC and a dramatic reduction in their expression in PDAC, consistent with the cysteine labeling patterns (Figure 4E). Despite this, the total cysteine pool in HCC was dramatically increased, suggesting HCC facilitates the accumulation of cysteine by other mechanisms (Figure S4A). In contrast, the total cysteine pool of PDAC was decreased (Figure S4B). These results demonstrate that transsulfuration capable tissues may maintain this capacity upon transformation, or may lose this capacity entirely.

**Figure 4.**
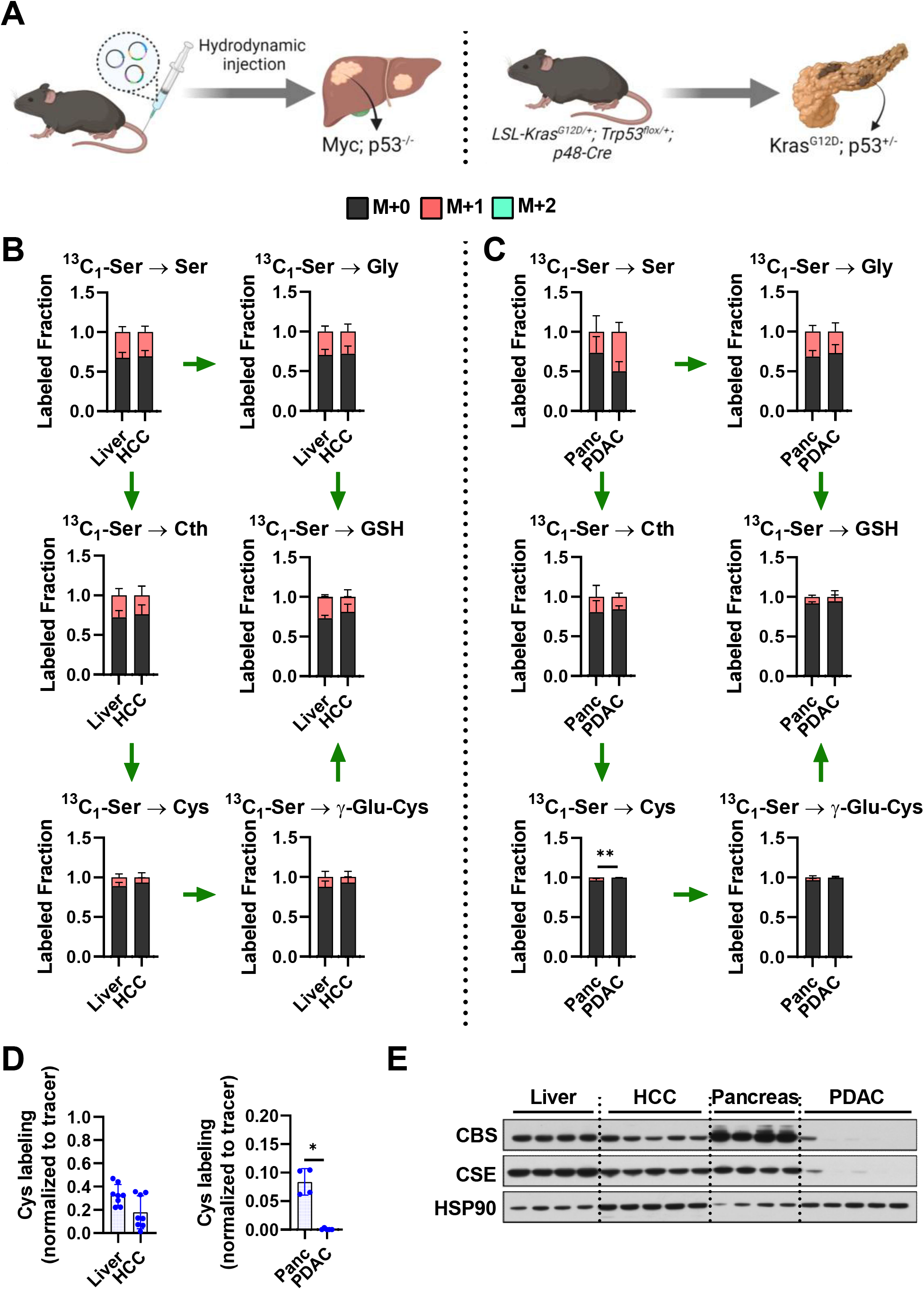
Tumorigenesis of Liver and Pancreas Induces Downregulation of De Novo Cysteine Synthesis. (A) Schematic for the generation of Myc; p53^-/-^ HCC and Kras^G12D^; p53^+/-^ for PDAC GEMM tumors. (B) Analysis of the fraction labeling in serine, glycine, cystathionine, glutathione, cysteine and γ-glutamylcysteine in control liver (N=8) and HCC tumors (N=8) following infusion with 1-[^13^C_3_]^-^ serine. (C) Analysis of the fraction labeling in serine, glycine, cystathionine, glutathione, cysteine and γ-glutamylcysteine in control pancreas (N=5) and PDAC tumors (N=5) following infusion with 1-[^13^C_3_]^-^ serine. (D) Fractional contribution of serine to intracellular cysteine synthesis in HCC and PDAC. Cysteine labeling was normalized to the fraction labeling of serine in each tissue. One healthy pancreas sample was excluded due to a division error. For B-D, data are presented as mean ± SD. (E) Immunoblots of cystathionine β-synthase (CBS) and cystathionine γ-lyase (CSE) for each tissue. HSP90 was used for the loading control. *p < 0.05, **p < 0.01, ***p < 0.001, and ****p < 0.0001. Ser, serine; Cth, cystathionine; Cys, cysteine; Gly, glycine; GSH, glutathione; γ-Glu-Cys, γ-glutamylcysteine

### De Novo Cysteine Synthesis Does Not Contribute to the Cysteine Pool of Lung Tumors

We next wanted to explore whether a transsulfuration incapable tissue could gain the use of this pathway upon transformation. To this end, we explored the transsulfuration capacity of lung tumors. Thus, we employed 1-[^13^C_1_]-serine tracing using three different GEM models (GEMMs). Given the role of NRF2 in cysteine metabolism, *LSL-Kras^G12D/+^; Trp53^flox/flox^* (KP) mice and *LSL-Kras^G12D/+^; Trp53^flox/flox^; LSL-Nfe2l2^D29H^* (KPN) mice (both C57BL/6J background) were used for lung adenocarcinoma (LUAD) (Figure 5A). We also examined the activity of the transsulfuration pathway in small cell lung cancer, given the very different cell of origin of this lung cancer cell type from non-small cell lung cancer. For the SCLC model, *Rb1^flox/flox^; Trp53^flox/fox^; Myc^LSL/+^ or Rb1^flox/flox^; Trp53^flox/fox^; Myc^LSL/LSL^* (RPM(M)) mice (mixed C57BL/6/FVB/129 background) were used (Figure 5A). In LUAD, the cystathionine labeling was similar to control lung, which was not influenced by Nrf2 mutation (Figure 5B). In contrast, SCLC demonstrated a significant reduction of cystathionine labeling compared with control (Figure 5C). Most importantly, none of the GEM tumors demonstrated cysteine labeling from serine, indicating a lack of transsulfuration of serine to cysteine (Figure 5B-D). Despite a lack of de novo cysteine synthesis, all tumor models accumulated tumoral cysteine, suggesting other mechanisms of cysteine accumulation (Figure 5D, S5A, and S5B). Immunoblotting for CBS and CSE revealed that both LUAD models (KP and KPN models) had down-regulated CBS compared to normal lung tissue, but CSE was overexpressed (Figure 5E). In contrast, the patterns in SCLC tumors were reversed, with a modest increase in CBS and downregulation of CSE (Figure 5E). These results demonstrate that lung tumors do not acquire de novo cysteine synthesis capacity and accumulate cysteine via other mechanisms.

**Figure 5.**
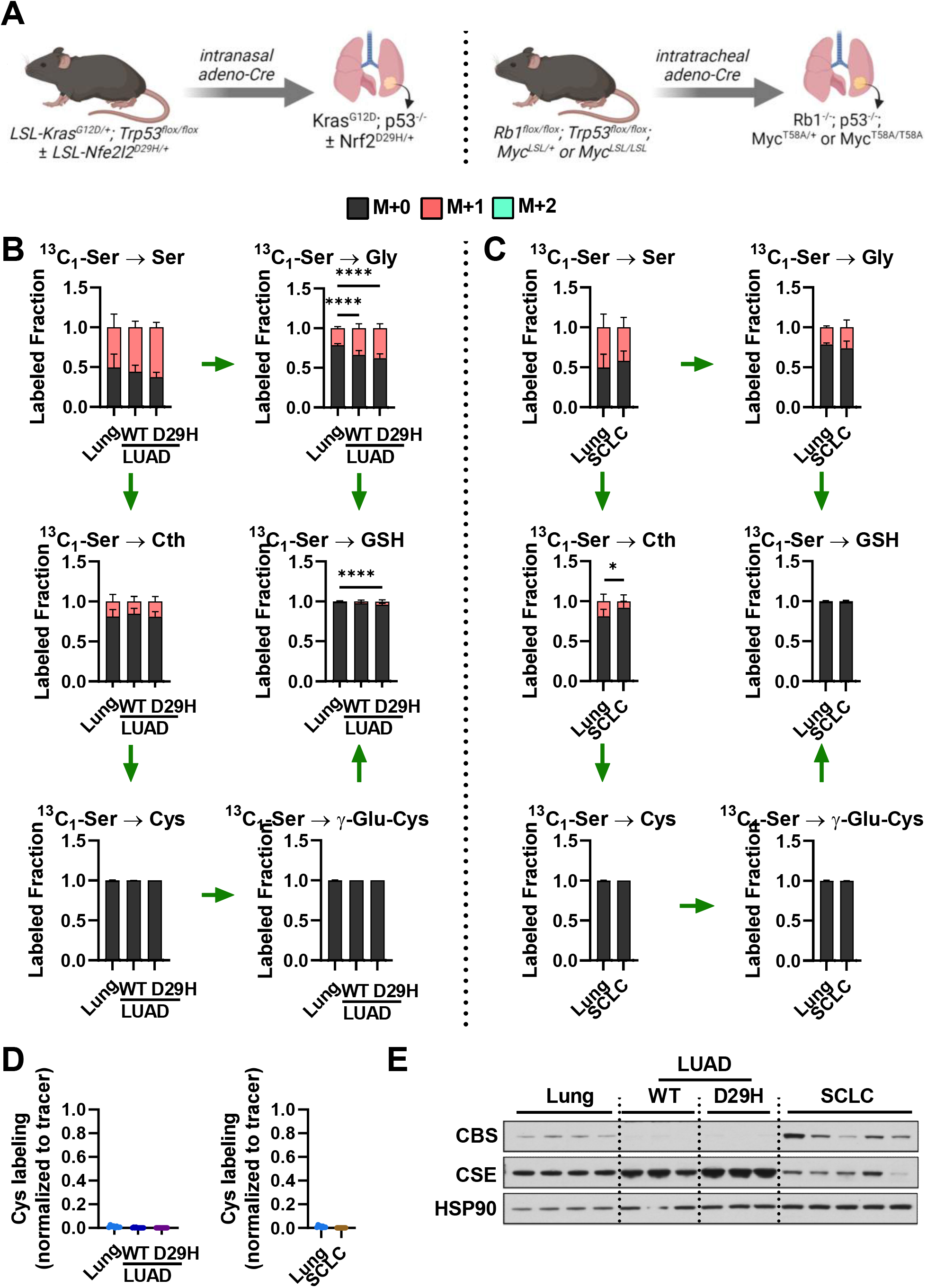
De Novo Cysteine Synthesis Does Not Contribute to the Cysteine Pool of Lung Tumors. (A) Schematic for the generation of Kras^G12D^; p53^-/-^ and Kras^G12D^; p53^-/-^; Nrf2^D29H^ LUAD, and Rb1^-/-^; p53^-/-^; Myc^T58A/+^ or Myc^T58A/T58A^ SCLC GEMM tumors. (B) Analysis of the fraction labeling in serine, glycine, cystathionine, glutathione, cysteine and γ-glutamylcysteine in control lung (N=8) and Nrf2^WT^ lung adenocarcinoma (N=10) and Nrf2^D29H^ lung adenocarcinoma (N=10) following infusion with 1-[^13^C_3_]^-^ serine. (C) Analysis of the fraction labeling in serine, glycine, cystathionine, glutathione, cysteine and γ-glutamylcysteine in in control lung (N=8) and small cell lung cancer (SCLC, N=10) following infusion with 1-[^13^C_3_]^-^ serine. N.B. the control lung samples in (C) are the same as in (B). (D) Fractional contribution of serine to intracellular cysteine synthesis in LUAD and SCLC. Cysteine labeling was normalized to the fraction labeling of serine in each tissue. For B-D, data are presented as mean ± SD. (E) Immunoblots of cystathionine β-synthase (CBS) and cystathionine γ-lyase (CSE) for each tissue. HSP90 was used for the loading control. *p < 0.05, **p < 0.01, ***p < 0.001, and ****p < 0.0001. Ser, serine; Cth, cystathionine; Cys, cysteine; Gly, glycine; GSH, glutathione; γ-Glu-Cys, γ-glutamylcysteine

### Cystine is a Major Contributor to the Cysteine Pool in Tumors

The accumulation of cysteine in the lung and HCC models suggests that these tumors may be fueling their cysteine pool from exogenous cyst(e)ine. To interrogate the contribution of cystine to the cysteine pool and downstream metabolites, we performed ^13^C_6_-cystine tracing in the HCC, PDAC, and LUAD GEMMs and the respective normal controls. We observed significant labeling in the cysteine pool in tumors, with higher labeling in cysteine synthesis deficient tumors (50%) compared to cysteine synthesis competent HCC (19%) (Figure 6A). HCC cysteine labeling was not significantly different from normal liver, while PDAC labeling was higher than pancreas and LUAD labeling was lower than normal lung (Figure 6A). Moreover, Nrf2^D29H^ increased cysteine labeling in tumors (Figure 6A). γ-glutamylcysteine labeling patterns mirrored cysteine labeling, apart from in lung tumors, where labeling was dramatically lower. Despite this, lung tumors had higher glutathione levels and labeling, with Nrf2^D29H^ tumors demonstrating significantly higher levels and labeling than Nrf2^WT^ tumors (Figure 6B and 6C). The incongruence between γ-glutamylcysteine labeling and GSH labeling in the lung tumors suggests potential dilution of the γ-glutamylcysteine pool by another source or cell type. HCC also demonstrated increased GSH labeling compared to normal liver, while PDAC demonstrated significantly lower labeling (Figure 6C). Despite this, HCC demonstrated lower GSH levels, and PDAC demonstrated higher GSH levels (Figure 6B). Immunoblotting revealed very low xCT expression in HCC, like normal liver, with similar expression of GCLC, GCLM and GSS between HCC and normal liver (Figure 6D). Normal pancreas and PDAC had high expression of xCT, with PDAC upregulating GCLC and GCLM, despite a reduction in GSH labeling (Figure 6D). LUAD upregulated xCT and GCLC, with Nrf2 promoting a further increase in xCT, GCLC and GCLM (Figure 6D). These findings reveal that cystine is a major contributor to the cysteine pool in tumors, but glutathione metabolism displays complex regulation across diverse tumor types.

**Figure 6.**
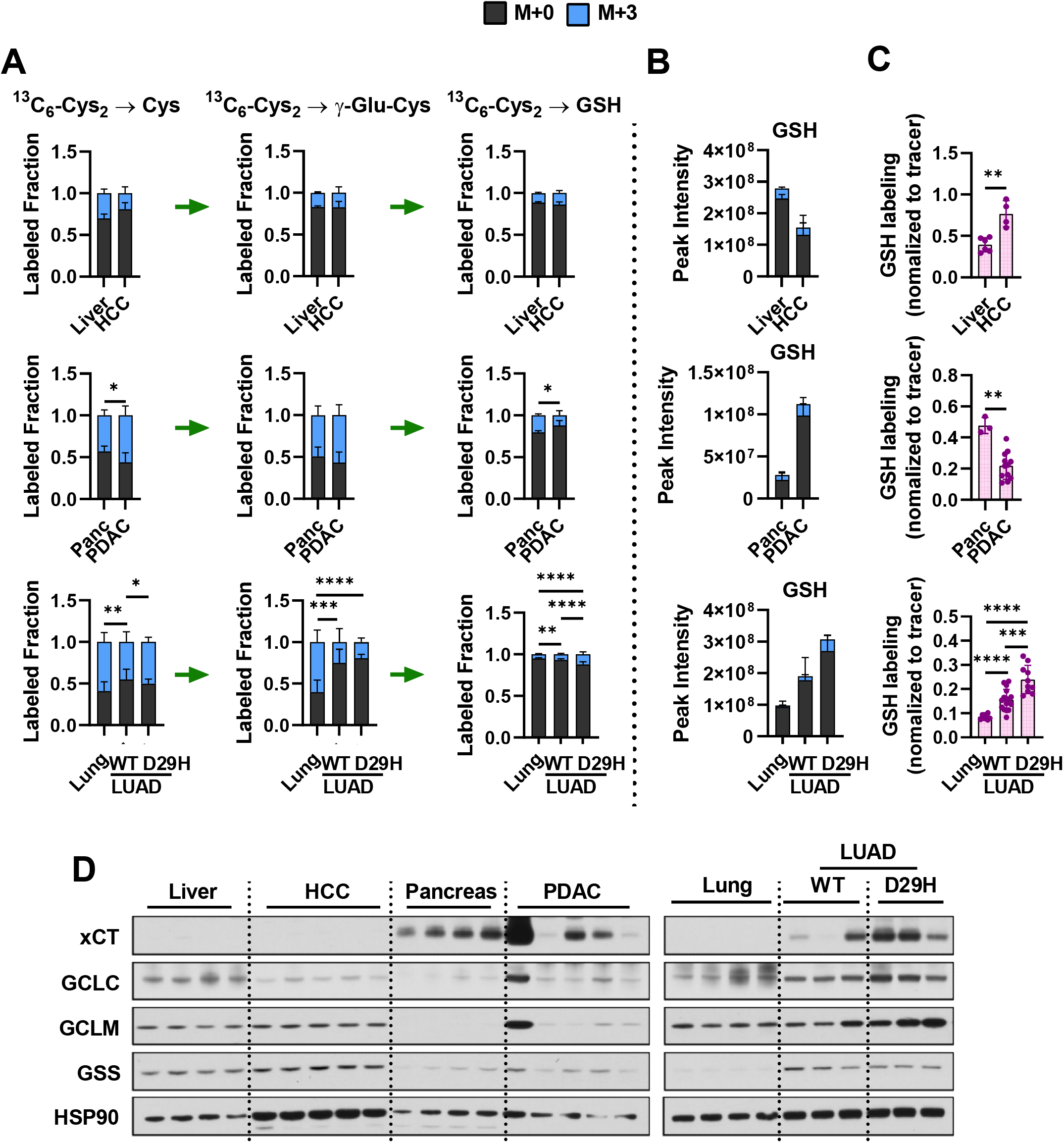
Cystine is a Major Contributor to the Cysteine Pool in Tumors. (A) Analysis of the fraction labeling in cysteine, γ-glutamylcysteine, and glutathione in control liver (N=6), HCC (N=4), control pancreas (N=3), PDAC (N=12), control lung (N=10), Nrf2^WT^ LUAD (N=16), and Nrf2^D29H^ LUAD (N=10) following infusion with ^13^C_6_-cystine. (B) Total signal of glutathione in the tissues from (A). (C) Fractional contribution of cystine to glutathione in each tissue from (A). Glutathione labeling was normalized to the fraction labeling of cysteine in each tissue. For (A-C), data are presented as mean ± SD. (D) Immunoblots of cystine/glutamate antiporter (xCT), glutamate-cysteine ligase catalytic subunit (GCLC), glutamate-cysteine ligase modifier subunit (GCLM), and glutathione synthetase (GSS) for each tissue. HSP90 was used for the loading control. *p < 0.05, **p < 0.01, ***p < 0.001, and ****p < 0.0001. Cys, cysteine; GSH, glutathione; γ-Glu-Cys, γ-glutamylcysteine.

### Cysteine Oxidation is Rewired in Tumors

In diverse human cancers, cysteine oxidation shifts toward enhancing adaptation of cancerous cells to reduce oxidative stress through increase of their antioxidative capacity (Chen et al., 2022). CDO1 catalyzes the rate limiting step in cysteine oxidation and is frequently downregulated by promoter methylation in many cancers (Brait et al., 2012). To examine cysteine oxidation in tumors, we analyzed the labeled fraction and total amount of cysteine, hypotaurine, and taurine (Figure 7A, 7B and S7A-S7E). We could not detect labeling in taurine except in normal liver (~ 6%) following 4 hours of [^13^C_6_]-cystine infusion (Figure 7A), so most analyses were limited to hypotaurine. When normalized to cysteine labeling, we found that HCC significantly upregulated hypotaurine synthesis compared to normal liver (Figure 7A-C). Examination CDO1 and CSAD protein expression revealed downregulation of CDO1, but upregulation of CSAD (Figure 7D). Hypotaurine can also be generated by CoA turnover and ADO, which mediates cysteamine metabolism to hypotaurine, was also increased (Figure 7D). Unfortunately, we did not detect pathway specific metabolites that can distinguish the CSAD vs ADO pathways, but CoA turnover is known to be very slow (Orsatti et al., 2021), suggesting label may be coming via the CDO1 pathway. Despite a reduction in CDO1, cysteine levels were significantly higher in HCC (Figure 7B), which may mediate increased entry of cysteine into the CDO1 pathway regardless of CDO1 expression level. By contrast, we observed that PDAC had significantly lower hypotaurine synthesis, consistent with both a decrease in cysteine availability and lower expression of both CDO1 and CSAD compared with normal pancreas tissues (Figure 7A-D and S7C). Interestingly, PDAC showed overexpression of ADO (Figure 7D). It is important to note that because ADO also has a role as a protein cysteine dioxygenase (Masson et al., 2019), its expression in tumors may be independent of taurine metabolism. In contrast with both HCC and PDAC, cysteine oxidation in LUAD was relatively insensitive to tumorigenesis (Figure 7B-7D). Finally, while we could not draw conclusions about taurine synthesis in most tissues due to limited labeling in the 4-hour period, we observed a reduction in labeling in the liver consistent with FMO1 downregulation (Figure 7D). FMO1 downregulation was also observed in LUAD tumors, although the impact on taurine labeling could not be evaluated (Figure 7D). Interestingly, PDAC demonstrated FMO1 overexpression, consistent with an increase in total taurine levels compared to normal pancreas (Figure 7D, and S7D). Because hypotaurine labeling was reduced in PDAC (Figure 7C), taurine accumulation is likely to come from exogenous sources. These findings demonstrate that tumors rewire cysteine oxidation and some tumors may accumulate taurine via alternative mechanisms.

**Figure 7.**
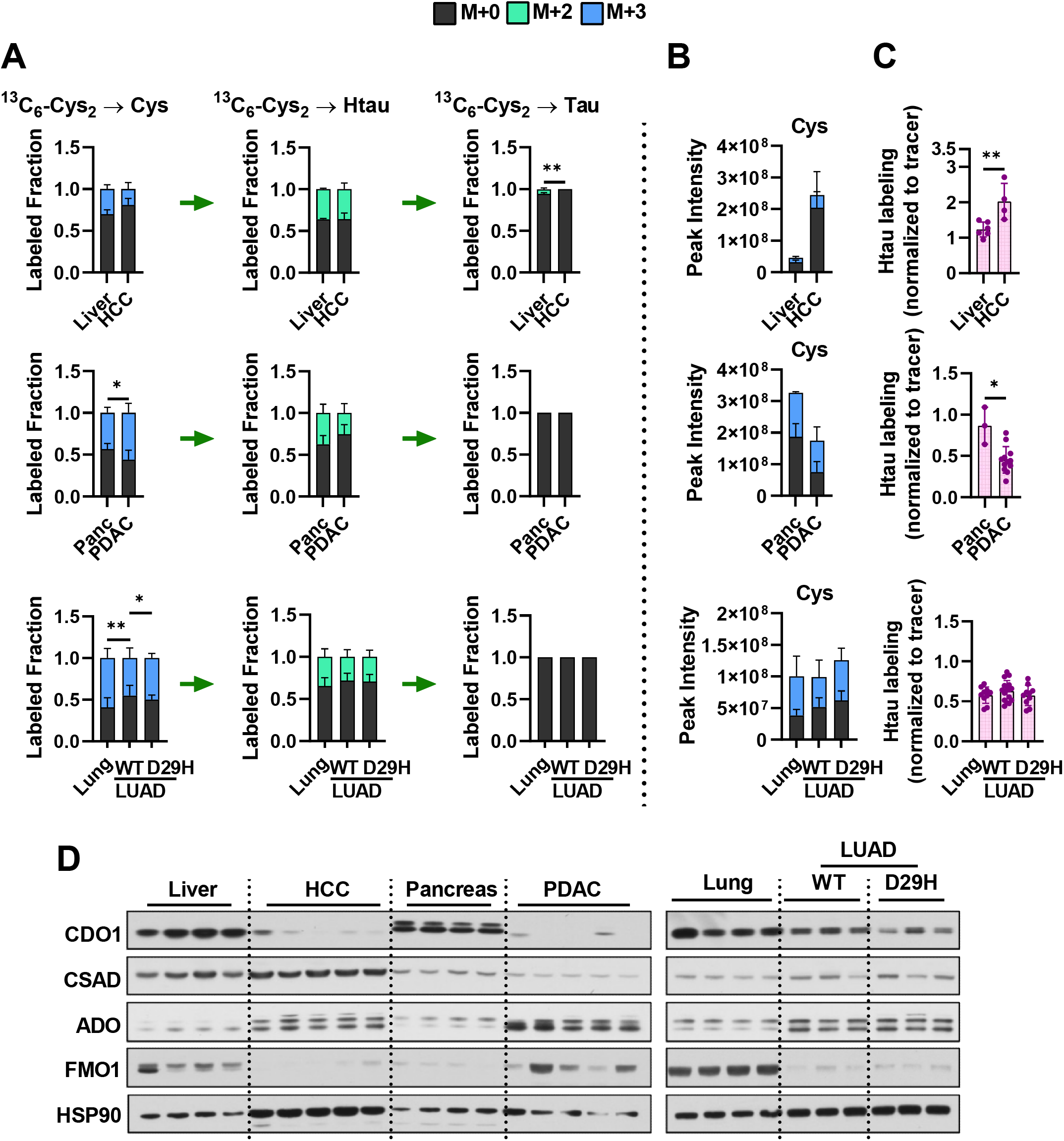
Cysteine Oxidation is Rewired in Tumors. (A) Analysis of the fraction labeling in cysteine, hypotaurine, and taurine in control liver (N=6), HCC (N=4), control pancreas (N=3), PDAC (N=12), control lung (N=10), Nrf2^WT^ LUAD (N=16), and Nrf2^D29H^ LUAD (N=10) following infusion with ^13^C_6_-cystine. (B) Total signal of cysteine in the tissues form (A). (C) Fractional contribution of cystine to hypotaurine in each tissue from (A). Hypotaurine labeling was normalized to the fraction labeling of cysteine in each tissue. For (A-C), data are presented as mean ± SD. (D) Immunoblots of cysteine dioxygenase type 1 (CDO1), cysteine sulfinate decarboxylase (CSAD), 2-aminoethanethiol (cysteamine) dioxygenase (ADO), and Flavin containing monoxygenase 1 (FMO1) for each tissue. HSP90 is used for the loading control. For (D), the same samples were analyzed as in Figure 6D and they share the same loading control (HSP90). *p < 0.05, **p < 0.01, ***p < 0.001, and ****p < 0.0001. Cys, cysteine; Htau, hypotaurine; Tau, taurine.

## DISCUSSION

In this study, we comprehensively evaluated the contribution of transsulfuration and exogenous cyst(e)ine to the cysteine pool and its downstream intermediates using ^13^C-based stable isotope tracers in vivo and in vitro. In both cultured cancer cell lines and tissues in vivo, we find that cystathionine is robustly labeled from serine, while cysteine labeling is slow or absent, even when exogenous cystine is removed from the culture system. CBS is considered the rate limiting enzyme of transsulfuration, and its activity is positively regulated by S-adenosylmethionine availability to promote entry of homocysteine into the transsulfuration pathway (Ascenção and Szabo, 2022). However, our results demonstrate that the second step of transsulfuration mediated by CSE is likely to restriction de novo cysteine synthesis in vitro as well as in vivo. Recently, interleukin 1 receptor accessory protein (IL1RAP) was identified as a novel positive regulator of cysteine availability that functions through both the regulation of xCT and transsulfuration (Zhang *et al*., 2021). Interestingly, IL1RAP promotes transsulfuration through the transcriptional regulation of CSE, suggesting CSE is the limiting component. Moreover, ATF4 transcriptionally regulates CSE (Bai et al., 2021), suggesting amino acid stress promotes cysteine synthesis at the CSE step. Therefore, it’s possible that the regulation of CBS is important to avoid the toxicity associated with homocysteine accumulation, while regulation of CSE is more important to control cysteine synthesis, but more work is needed.

Given the importance of cysteine, why is the transsulfuration pathway not a major contributor to the cysteine pool in tumors? We find that de novo cystine synthesis is either inactive (lung cancer) or downregulated (HCC and PDAC) during tumorigenesis. Downregulation of transsulfuration is associated with a decreased expression of both CBS and CSE. Prior studies have reported the downregulation of CBS in both HepG2 cells and HCC patients, which is associated with poor prognosis (Kim et al., 2009; Wang et al., 2018). Downregulation of CSE is also observed in HCC and associated with poor prognosis (Lin et al., 2021). The analysis of CBS and CSE expression in PDAC is complicated by the dense stroma typical of these tumors (Helms et al., 2020), which is comprised of fibroblasts, immune cells, and other cell types. While this stroma can account for half of the tumor cellularity, it is unlikely to completely account for the complete absence of cysteine synthesis and CBS/CSE expression we observe in the GEMMs. In contrast to HCC, low expression of CBS in PDAC is associated with better outcomes (Ascenção and Szabo, 2022). In addition to their role in cysteine synthesis, CBS and CSE play a role in hydrogen sulfide (H2S) generation, which can be toxic in high concentrations (Ascenção and Szabo, 2022). Thus, tumors in which adequate cysteine is supplied from other pathways may downregulate these enzymes to limit the other metabolic consequences of these enzymes. However, other cancer types not evaluated in our study, such as colorectal carcinoma, are reported to increase CBS expression, which is associated with worse outcomes. Additional studies are needed to evaluate CBS and CSE expression and cysteine synthesis activity in other models.

We find that in normal tissues, cyst(e)ine readily labels the cysteine pool. However, xCT expression alone is not a good predictor of cysteine labeling. The uptake of cystine and its reduction to cysteine via xCT is influenced by other factors, including both intracellular and extracellular glutamate availability, cellular reducing potential, and xCT post-translational modification and subcellular localization of xCT (Combs and DeNicola, 2019; Mukhopadhyay *et al*., 2021). Indeed, we find that concomitant with high xCT expression the brain tissues also have high glutamate levels, which likely limit cystine/glutamate exchange. Moreover, xCT knockout mice are viable (Sato et al., 2005), indicating that other cystine and/or cysteine transporters can support normal cysteine homeostasis. Cysteine transporters are poorly characterized, particularly in the context of cancer. Interestingly, we find that despite an increase in the total cysteine pool in HCC, this cannot be accounted for by an increase in transsulfuration or cystine uptake, suggesting that HCC tumors have an alternative source of cysteine. Glutathione degradation via gamma-glutamyl transpeptidase may locally generate available cysteine (Asantewaa and Harris, 2021), or tumors may recycle micropinocytosis-derived protein to contribute to the cysteine pool as has been shown in HCC cell lines (Byun et al., 2022). Additional work is needed to understand the reliance of HCC tumors on other cysteine sources. Given this potential metabolic flexibility, it may be challenging to target cysteine availability in HCC.

We examined the metabolism of cysteine to downstream metabolites in tumors. Cancer cells are generally thought to have an increased demand for antioxidant protection, particularly via glutathione synthesis (Pavlova *et al*., 2022). We find that both HCC and LUAD tumors show increased contribution of cysteine to the glutathione pool, with NRF2 activation further promoting glutathione synthesis as expected. Interestingly, PDAC decreased the contribution of cysteine to glutathione despite having a very high total glutathione content. PDAC is has high macropinocytic activity (Commisso et al., 2013), which facilitates the uptake of protein from the protein rich extracellular environment for degradation to supply the intracellular amino acid pools. Glutathione is about 7 times higher in PDAC interstitial fluid compared to plasma (184μM vs 26μM) in a GEMM (Sullivan et al., 2019), raising the interesting possibility that PDAC can acquire glutathione via micropinocytosis to supply the intracellular pool. In addition to glutathione synthesis, we examined cysteine oxidation to taurine as another downstream cysteine catabolic pathway. Taurine is thought to be predominantly synthesized in the liver, and to a lesser extent in other tissues, then released into circulation for uptake via the ubiquitously expressed taurine transporter (TAUT, SLC6A6) (Stipanuk, 2004a). However, our results suggest that kidney, liver and brain (including cerebellum) label their hypotaurine pool from circulating hypotaurine, suggesting uptake of hypotaurine itself in these tissues. Hypotaurine is transported by γ-aminobutyric acid transporter type 2 (GAT2, SLC6A13) and TAUT (Nishimura et al., 2018). GAT2 is primarily expressed in not only brain including cerebellum but also liver and kidney (Zhou and Danbolt, 2013). Moreover, pancreas expresses high levels of CDO1, but lacks the expression of FMO1, raising the possibility that pancreas synthesizes high levels of hypotaurine for export.

Interestingly, our findings demonstrate that all tumors show downregulation of CDO1 expression, consistent with the frequent epigenetic silencing of this gene in human cancer (Choi et al., 2017; Igarashi et al., 2017; Kojima et al., 2018; Maekawa et al., 2020; Nishizawa et al., 2019; Ooki et al., 2017; Yin et al., 2020). While we previously reported that Nrf2 promoted CDO1 accumulation in a Keap1^R554Q^ model of LUAD (Kang *et al*., 2019), this analysis was on the low-grade tumors that are most frequent in the Keap1^R554Q^ model. The Nrf2^D29H^ tumors of sufficient size for excision for metabolomics analysis in this study were instead higher-grade adenocarcinomas that we find downregulate Nrf2 and target protein expression (DeBlasi et al., 2022), which will influence CDO1 stabilization. We previously reported that the downregulation of taurine synthesis conserves both cysteine and NADPH (Kang *et al*., 2019), but taurine is still a requirement due to its role in osmolarity control, mitochondrial translation, redox regulation, and other processes (Ward and DeNicola, 2019). We find that FMO1 is overexpressed in PDAC compared to normal pancreas, which is correlated with a dramatic increase in taurine, suggesting that PDAC may uptake and metabolize hypotaurine to taurine. Consistently, we see an increased contribution of circulating hypotaurine to the hypotaurine pool in HCC, suggesting that tumors may generally supply their demands for taurine from the circulation. Finally, we find that ADO is generally overexpressed in tumors, consistent with its reported regulation by hypoxia (Masson *et al*., 2019). While ADO has a role in protein cysteine oxidation, it may also play a role in cysteine metabolism in tumors. Due to technical reasons, we could not detect cysteamine in our tissues, and therefore cannot conclude anything about the role of ADO in hypotaurine metabolism.

### Limitations of Study

There are several limitations of our study. First, we examined cysteine labeling and downstream metabolism at a single time point after a four-hour infusion with ^13^C-serine or ^13^C-cystine, which allowed us to examine the contribution of exogenous cyst(e)ine and the transulfuration pathway to the cysteine pool and downstream metabolism. We were unable to evaluate the contribution of other sources of cysteine, including glutathione, protein, and even circulating cysteine precursors like cystathionine, with this approach. We analyzed steady state labeling of metabolites in cysteine metabolic pathways but did not assay flux over time like what was recently reported for TCA cycle flux (Bartman et al., 2021), which would provide additional information. Our analyses were also limited to macrodissected tissues that are comprised of multiple cell types. Combining labeling with spatial metabolomics will be critical to deconvoluting where reactions are occurring (Wang et al., 2022). Finally, our studies are limited to mice and while sulfur metabolism is highly conserved, there may be microbiome, diet, and environmental differences that influence the translation of our studies to humans.

## Acknowledgements

We would like to thank all members of the DeNicola laboratory and Isaac Harris for their very helpful discussions. This work was supported by grants from the NIH/NCI R37CA230042 to G.M.D and P01CA250984 to G.M.D and E.R.F. This work was also supported by Miles for Moffitt funds awarded to the Lung Cancer Metabolism Working Group, and the Proteomics/Metabolomics Core, which is funded in part by Moffitt’s Cancer Center Support Grant (NCI, P30-CA076292). Biorender was used to generate figure schematics.

## Author Contributions

S.J.Y and G.M.D. designed the study and interpreted experimental results, S.J.Y and J.C. performed metabolomics, N.P-F performed western blots, H.D.A and E.R.F. provided SCLC mice, A.F and S.C. generated experimental animals and performed animal infusions, S.J.Y and G.M.D wrote the manuscript and all authors commented on it.

## Declaration of Interests

The authors declare no competing interests.

## METHODS

### KEY RESOURCES TABLE

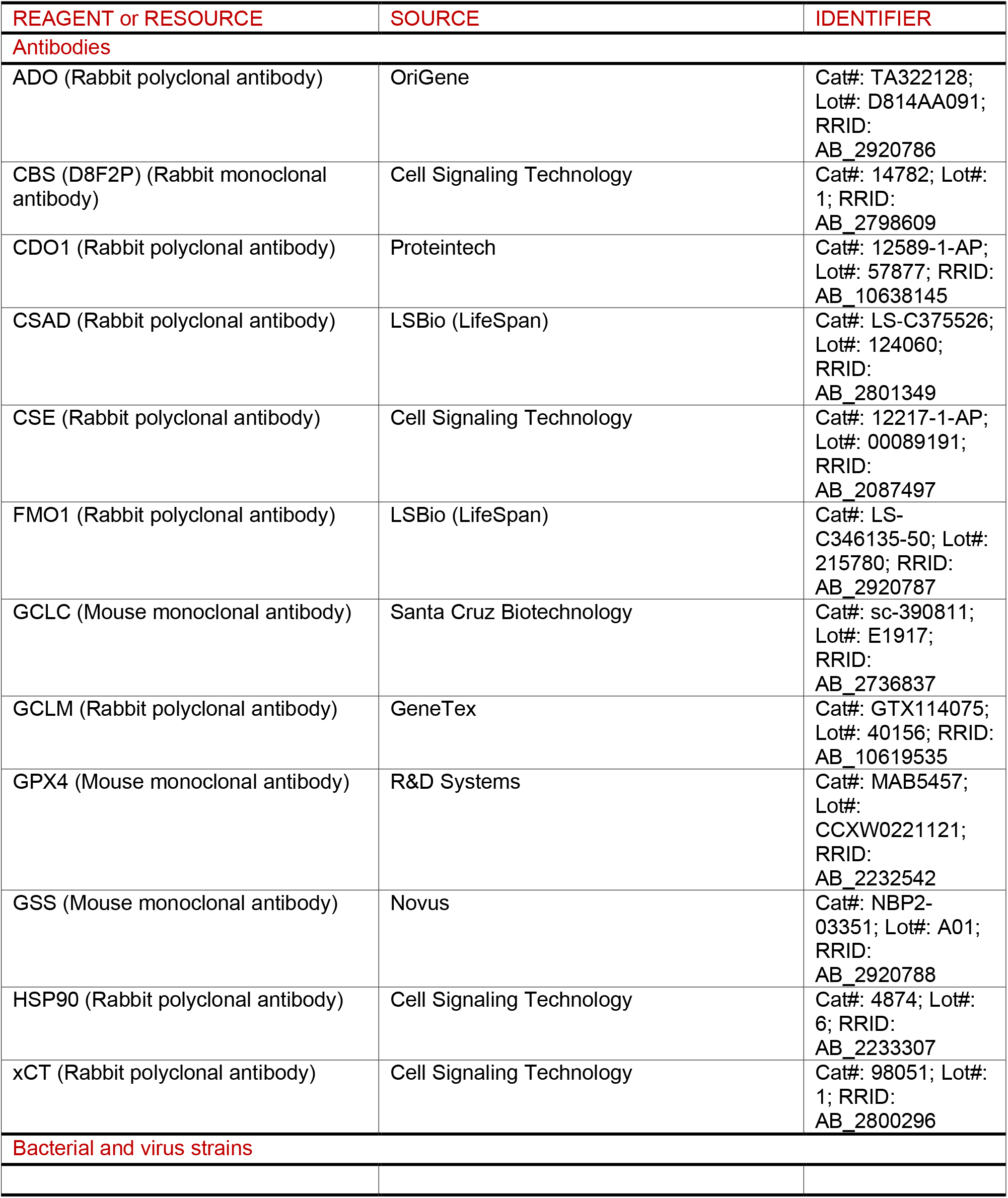
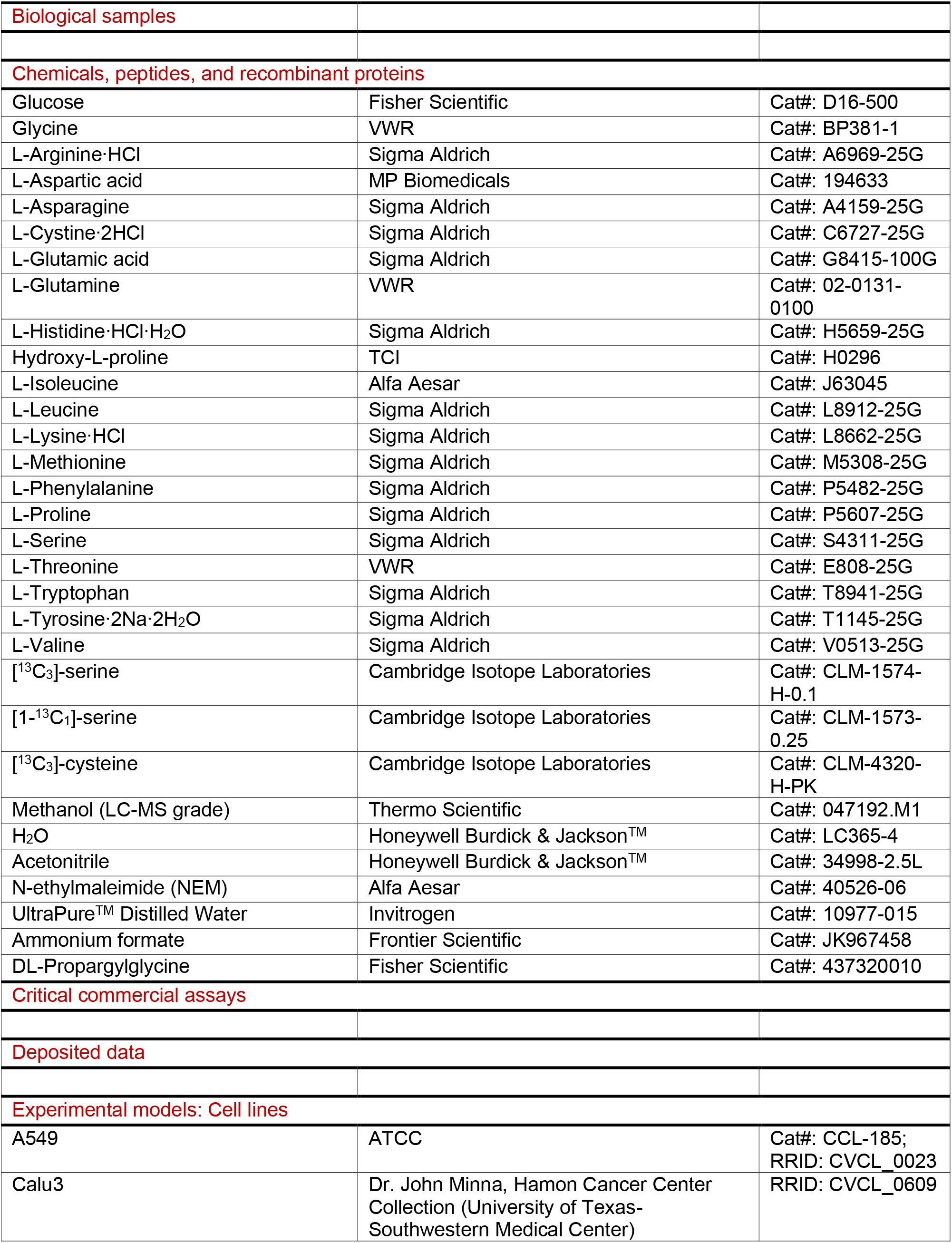
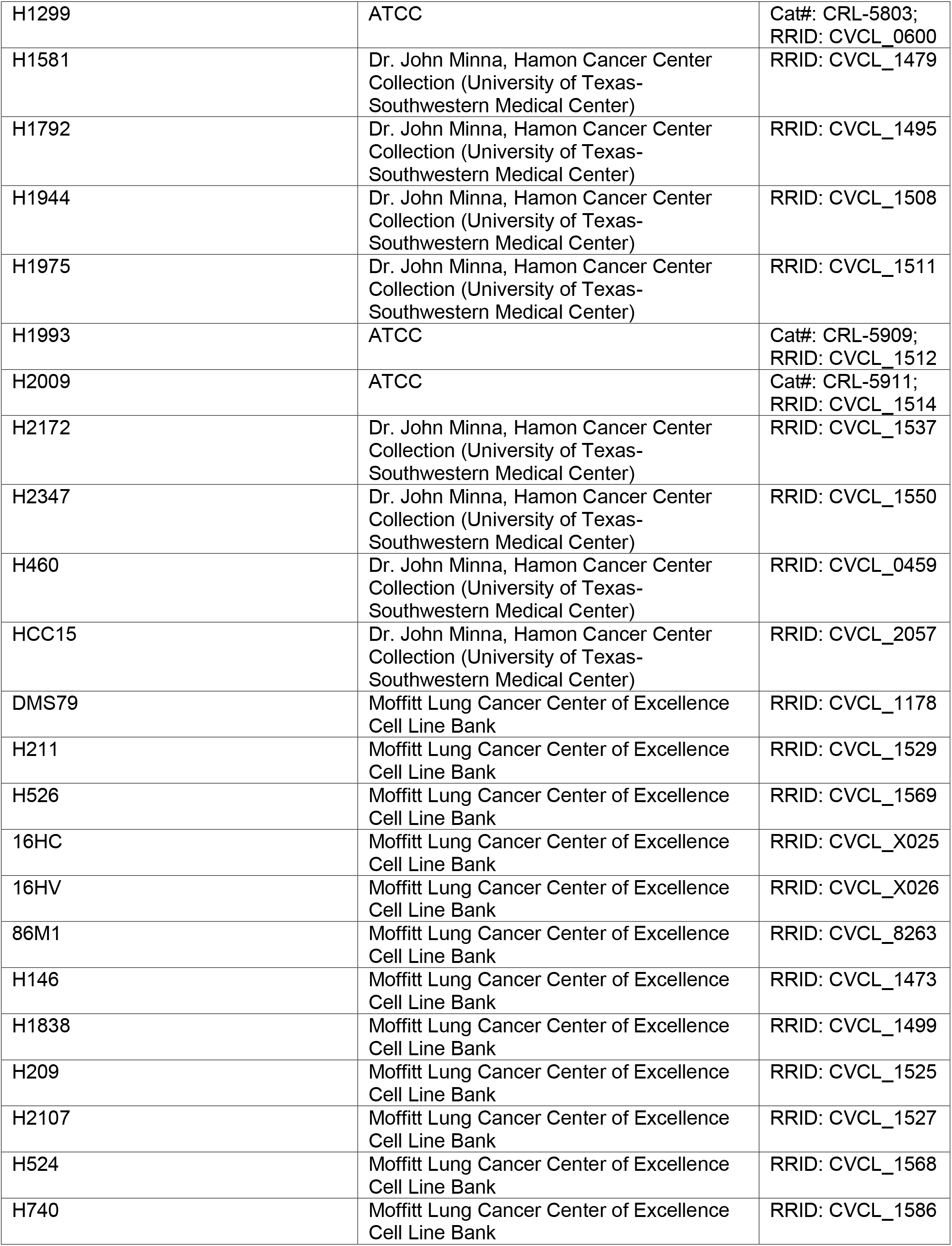
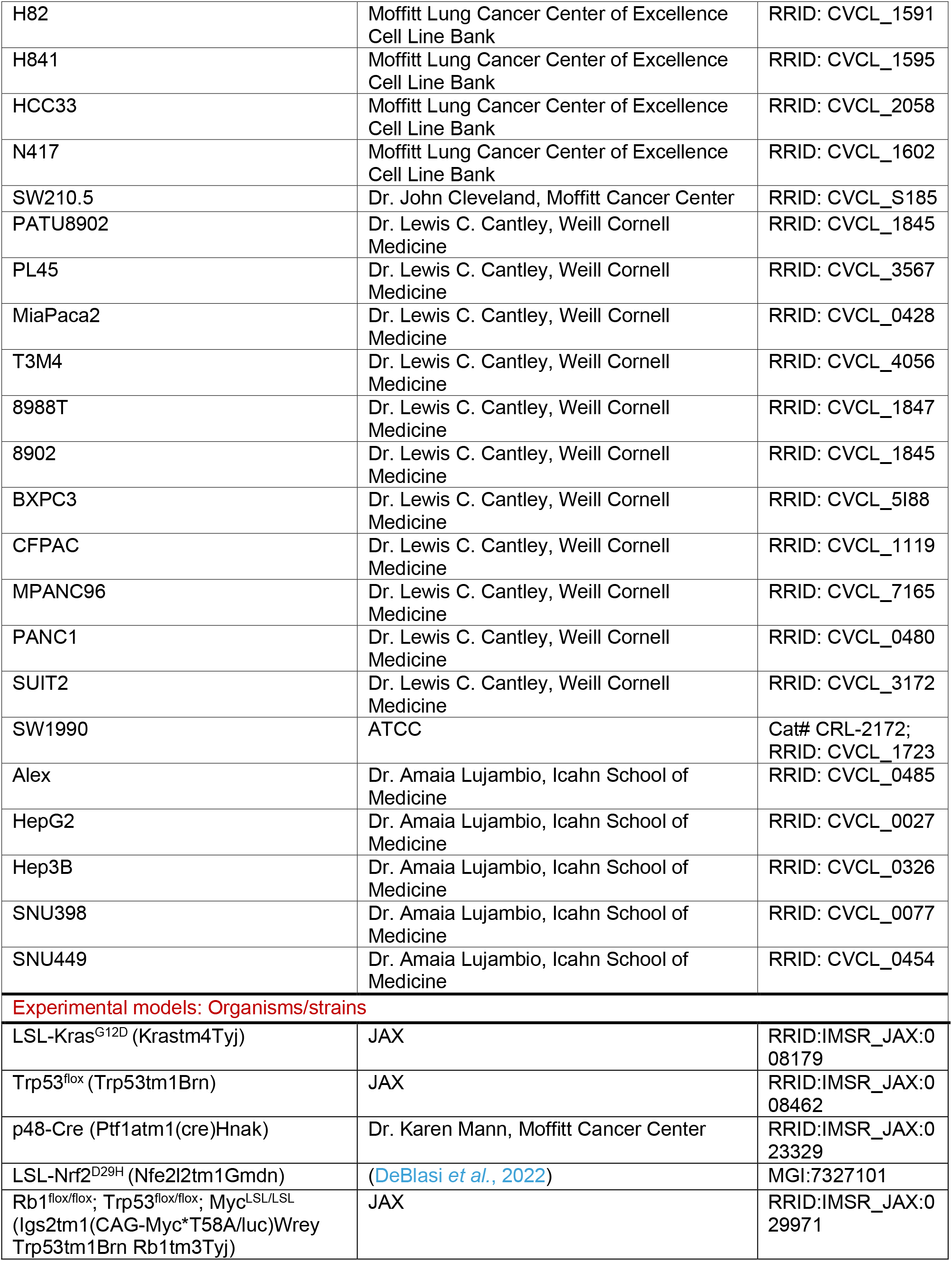
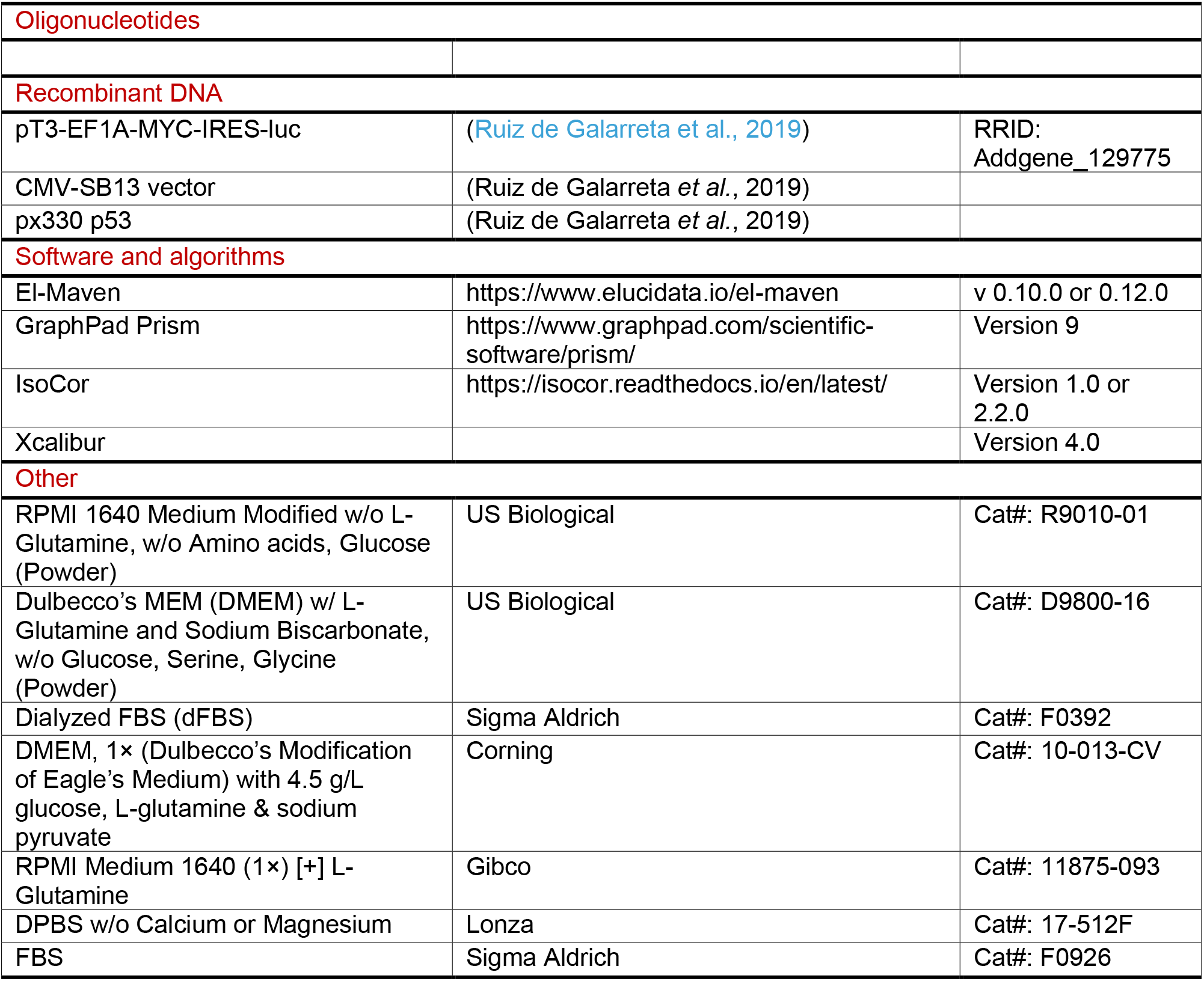

### RESOURCE AVAILABILITY

#### Lead Contact

Further information and requests for resources and reagents should be directed to and will be fulfilled by the Lead Contact, Dr. Gina M. DeNicola.

#### Material Availability

All unique reagents generated in this study are available from the Lead Contact without restriction.

#### Data and Code Availability

This study did not generate any unique datasets or code.

### EXPERIMENTAL MODEL AND SUBJECT DETAILS

#### Generation of Experimental Animals

All animal experiments were performed following IACUC approved procedures (Protocols IS00003893R, IS00006358R, IS00007922R, IS00008736R, and IS00010348R). To generate Myc; p53^-/-^ HCC tumors (Ruiz de Galarreta *et al*., 2019), DNA was delivered to the liver of 8 week old C57BL/6J female via hydrodynamic tail vein injection to concomitantly integrate a Myc transposon into the mouse genome and delete *Trp53*. A volume of sterile saline equal to 10% of their body weight containing 10 μg of Myc-Luciferase plasmid, 10 μg of Cas9/sgp53 plasmid, and 2.5 μg of SB13 transposase plasmid was injected into the mouse tail vein. Liver tumors developed approximately 4-6 weeks later, which was monitored by bioluminescence imaging. To generate experimental PDAC mice, mice harboring *LSL-Kras^G12D^, Trp53^flox^* and *p48-cre* alleles were intercrossed to generate *LSL-Kras^G12D/+^; Trp53^flox/+^; p48-cre* experimental animals. Mice developed tumors with a median survival of approximately 6 months (Morton et al., 2010), and tumor development was monitored by abdominal palpation, followed by confirmation by ultrasound. To generate experimental animals with LUAD tumors, *LSL-Kras^G12D/+^, Trp53^flox^* and *LSL-Nrf2^D29H^* mice were intercrossed to generate *LSL-Kras^G12D/+^; Trp53^flox/flox^* and *LSL-Kras^G12D/+^; Trp53^flox/flox^; LSL-Nrf2^D29H^* mice. Mice were infected intranasally with 2×10^7^ PFU adenovirus (Ad5CMVCre, University of Iowa) under isofluorane anesthesia to initiate tumor development. Mice developed tumors with a median survival of approximately 4 months post infection as previously reported (DeBlasi *et al*., 2022; Jackson et al., 2005), and mice were used for experiments between 3-3.5 months when they displayed evidence of tumor burden such as rapid respiration. To generate experimental animals with SCLC tumors, *Rb1^flox/flox^; Trp53^flox/flox^; Myc^LSL/LSL^* mice were intercrossed to generate *Rb1^flox/flox^; Trp53^flox/flox^; Myc^LSL/+^* (RPM) and *Rb1^flox/flox^; Trp53^flox/flox^; Myc^LSL/LSL^* (RPMM) experimental animals (Mollaoglu et al., 2017). Mice were infected intratracheally with 7.5×10^7^ PFU adenovirus (Ad5CGRPCre, University of Iowa) under ketamine (100mg/kg)/xylazine (10mg/kg) anesthesia to initiate tumor development. Mice developed tumors around 9-10 weeks post infection, and tumor development was monitored by MRI.

#### Stable Isotope Animals Infusions

[^13^C_6_]-cystine was generated from [^13^C_3_]-cysteine by oxidation with H_2_O_2_. A 20mg/mL solution of [^13^C_3_]-cysteine was dissolved in sterile saline, followed by the addition of an equimolar volume of 30% H_2_O_2_ (0.94μL/mg cysteine) and cysteine was allowed to oxidize for 30 minutes at room temperature on a rocker, during which time the resulting cystine precipitated. The H_2_O_2_ was inactivated by heating at 60°C for 5 minutes. Cystine was resolubilized by the addition of 6N HCl. For infusion of stable isotope tracers, catheters were surgically implanted into the jugular vein 2-7 days prior to infusion and mice were allowed to recover prior to infusion. On day of experiment, catheters were connected to a syringe pump on a tether and swivel system (SAI Technologies) to allow mice to freely move around the cage during infusions. A mouse harness with a spring prevented the tubing from disconnecting from the catheter. Syringes prefilled with saline containing 20 mg/mL of [1-^13^C_1_]-serine or [^13^C_6_]-cystine were loaded into the pump and tracers were infused at a rate of 120 μL/h for 4 h. During the final minutes of the infusion, blood was collected from the (cheek vein). Mice were sacrificed by cervical dislocation and organs of interest rapidly collected in cryovials and frozen in liquid nitrogen.

#### Cell Viability Test after Homocysteine and Cystathionine Rescue

Cystine free media was prepared from RPMI 1640 Medium powder lacking amino acids according to instructions and amino acids were added following the RPMI 1640 formulation except cystine. NSCLC cell lines (Calu3, H1944, H2009, H2347, and H1792) were plated RPMI 1640 (5% FBS) in 96 well plate at density of 10,000 cells/well in a 100 μL final volume. The following day, the medium was aspirated and cells were washed with DPBS, followed by with 100 μL of RPMI (5% dFBS) containing 0μM cystine or 200μM cystine. For rescue experiments, homocysteine or cystathionine was added into media for final concentration of 1 mM. For CSE inhibition experiments, propargylglycine (PPG) was added to the media for final concentration of 100 μM. Three days later, cells were fixed in iced cold 4% of paraformaldehyde at 4°C for 20 min, then stained with 50 μL of 0.1% crystal violet in 20% methanol on orbital shaker (room temperature, 30 min). The plate was washed twice with dH_2_O and dried. Crystal violet was solubilized in 100 μL of 10% acetic and the absorbance was read at 600 nm.

#### Stable Isotope Labeling in Cell Culture

To prepare medium including [^13^C_3_]-serine, RPMI 1640 Medium powder without glucose and amino acids (US Biological) and DMEM powder without glucose, glycine, and serine (US Biological) were reconstituted following manufacturer’s instructions. Glucose and amino acids were added to match the RPMI 1640 and DMEM formulation except serine. RPMI feeding media contained 300 μM [^13^C_3_]-serine + 5% dFBS and DMEM contained 400 μM [^13^C_3_]-serine + 10% dFBS. Both were supplemented with 1% Pen/Strep. Cell lines were plated in 6 well plates so they were 70% confluent at the time of extraction. Cells were preconditioned in medium including dFBS overnight (RPMI with 5% dFBS for NSCLC and SCLC cell lines; DMEM with 10% dFBS for PDAC and HCC cell lines). Prior to labeling, the cells were washed with 1 mL serine-free medium and then fed with medium containing [^13^C_3_]-serine for 4 hours prior to extraction.

#### Metabolomics Sample Preparation

Cells were washed with ice-cold DPBS, followed by aspiration of medium. 500 μL of ice-cold extraction solvent (80% methanol and 20% water including was 10 mM ammonium formate and 25 mM NEM, pH 7.0) was added to each well. After 30 min of incubation at 4°C, cells were scraped and, the supernatant moved to a 1.5mL tube and the debris cleared by centrifugation (17,000 g, 4°C, 20 min). Extracts were stored at −80°C until analysis. Cell numbers were counted by Scepter 2.0 cell counter and used to calculate intracellular metabolite concentrations. To extract metabolites from tissues, the frozen tissues were pulverized with a pre-chilled Bio-Pulverizer (59012MS, BioSpec). After weighing the tissues, the extraction solvent (80% methanol and 20% water including was 10 mM ammonium formate and 25 mM NEM, pH 7.0) was added to the pulverized tissue for a final concentration of 50 mg tissue/mL extraction solvent for 30 min at 4°C. To extract metabolites from serum, 390 μL of extraction solvent (82% methanol and 18% water including was 10 mM ammonium formate and 25 mM NEM, pH 7.0) was added to 10 μL of serum, followed by incubation at −80°C for 30 min. Debris was cleared by centrifugation (17,000 g, 4°C, 20 min). Extracts were stored at −80°C until analysis.

#### LC-MS Analysis and Data Processing

The instrumental conditions of LC-MS analysis were optimized based on previously established methods (Kang *et al*., 2021). The chromatography system for separation was the Vanquish UPLC system equipped with a SeQuant ZIC-pHILIC column (150 × 4.6 mm, 5 μm, MilliporeSigma, Burlington, MA) connected to a SeQuant ZIC-pHILIC guard column (20 × 4.6 mm, 5 μm, MilliporeSigma, Burlington, MA) or an Atlantis Premier BEH Z-HILIC VanGuard FIT column (2.1 mm x 150 mm, 2.5 μm, Waters, Milford, MA). The column was kept in a 30°C column chamber 5 μL of sample loaded via auto-sampler. For the gradient, mobile phase A (10 mM ammonium carbonate and 0.05% ammonium hydroxide in water) and mobile phase B (100% acetonitrile) were used as follow: 0 min, 20% of B; 13 min, 80% of B; 15 min, 20% of B; 20 min, 20% of B. For separated metabolite detection, a Q Exactive™ HF (QE-HF) Orbitrap mass spectrometer (Thermo Scientific, Waltham, MA) with H-ESI was used. The ions were detected by positive mode and the MS^1^ scan range was *m/z* 65-950. The capillary temperature and voltage were 30°C and 3.5 kV, respectively. The mass resolution was 120,000 and the AGC target was 3×10^6^. After data conversion from .raw to .cdf using Xcalibur (Version 4.0), further data processing for targeted metabolomics was performed by El-Maven (Version 0.10.0 or 0.12.0) and the default parameters were used for data processing except as follows: ionization mode, positive; Isotopic tracer, C_1_3; extracted-ion chromatogram (EIC) extraction window (+/-), 10.00 ppm. Identification of metabolites was performed based on retention time and exact precursor ion *m/z* in previously established authentic standard-based in-house library (Kang *et al*., 2021). The peak intensity of each EIC was measured as AreaTop (mean of three top points in the peak). For isotope correction, the extracted metabolite signals from El-Maven were loaded into the IsoCor (https://isocor.readthedocs.io/en/latest/; Version 1.0 or 2.2.0) as .tsv file according to the recommended format and processed with following parameters: Isotopic tracer, ^13^C; ‘Low resolution’ was selected; ‘Correct natural abondance of the tracer element’ was selected; Isotopic purity of the tracer, ^12^C was 0.01 and ^13^C was 0.99.

#### Immunoblotting

To prepare cell lysates, cells were washed with ice cold DPBS, detached from 6 well plates by scraping, transferred to microcentrifuge tubes, and pelleted. Cell pellets were lysed in RIPA lysis buffer (20 mM Tris-HCl, pH7.5; 150 mM NaCl, 1 mM EDTA, 1 mM EGTA, 1% NP-40, 1% sodium deoxycholate) containing protease inhibitors and phosphatase inhibitors on ice for 30 minutes. To extract protein from tissue samples, 25 μL of RIPA containing protease inhibitors and phosphatase inhibitors was added per 1 mg of tissue. After homogenization with a dounce homogenizer, the samples were incubated on ice for 30 minutes, followed by sonication for 5 minutes on medium power (30 seconds on/ 30 seconds off). Debris was cleared by centrifugation (13,000 g, 4°C, 15 min) and supernatants were stored at −20°C until analysis.

Protein quantification was conducted using DC assay (Bio-Rad) according to the manufacturer’s instructions. The protein lysates were combined with 6× loading buffer containing 2-mercaptoethanol and loaded on NuPAGE 4%-12% Bis-Tris Midi gels (Invitrogen). After separation of protein by SDS-PAGE, the proteins were transferred to 0.45 μm Nitrocellulose blotting membrane (Cytiva). Following the blocking of membrane with 5% non-fat milk in TBST for 30 minutes, the membranes were washed three times with TBST for 10 minutes each, and the membranes were incubated in primary antibodies at a 1:1000 dilution in 2% milk overnight at 4°C. After incubation, the membranes were washed again in TBST and developed with ECL using X-ray film. When comparing cell line or tissue samples across multiple membranes, the same lysate was loaded on multiple membranes to ensure exposures were equal.

### QUANTIFICATION AND STATISTICAL ANALYSIS

GraphPad Prism 9 was used for all statistical analysis. The Mann-Whitney test (nonparametric t-test) was conducted for statistical comparisons.

## Notes

### Competing Interest Statement

The authors have declared no competing interest.

## REFERNCES

Anderson, M.E. (1998). Glutathione: an overview of biosynthesis and modulation. Chem Biol Interact 111–112, 1–14. 10.1016/s0009-2797(97)00146-4.

Asantewaa, G., and Harris, I.S. (2021). Glutathione and its precursors in cancer. Curr Opin Biotechnol 68, 292–299. 10.1016/j.copbio.2021.03.001.

Ascenção, K., and Szabo, C. (2022). Emerging roles of cystathionine β-synthase in various forms of cancer. Redox Biol 53, 102331. 10.1016/j.redox.2022.102331.

Badgley, M.A., Kremer, D.M., Maurer, H.C., DelGiorno, K.E., Lee, H.J., Purohit, V., Sagalovskiy, I.R., Ma, A., Kapilian, J., Firl, C.E.M., et al. (2020). Cysteine depletion induces pancreatic tumor ferroptosis in mice. Science 368, 85–89. 10.1126/science.aaw9872.

Bai, X., Ni, J., Beretov, J., Wasinger, V.C., Wang, S., Zhu, Y., Graham, P., and Li, Y. (2021). Activation of the eIF2α/ATF4 axis drives triple-negative breast cancer radioresistance by promoting glutathione biosynthesis. Redox Biol 43, 101993. 10.1016/j.redox.2021.101993.

Bartman, C.R., Shen, Y., Lee, W.D., TeSlaa, T., Jankowski, C.S.R., Wang, L., Yang, L., Roichman, A., Bhatt, V., Lan, T., et al. (2021). Slow TCA flux implies low ATP production in tumors. bioRxiv, 2021.2010.2004.463108. 10.1101/2021.10.04.463108.

Brait, M., Ling, S., Nagpal, J.K., Chang, X., Park, H.L., Lee, J., Okamura, J., Yamashita, K., Sidransky, D., and Kim, M.S. (2012). Cysteine dioxygenase 1 is a tumor suppressor gene silenced by promoter methylation in multiple human cancers. PLoS One 7, e44951. 10.1371/journal.pone.0044951.

Byun, J.K., Lee, S., Kang, G.W., Lee, Y.R., Park, S.Y., Song, I.S., Yun, J.W., Lee, J., Choi, Y.K., and Park, K.G. (2022). Macropinocytosis is an alternative pathway of cysteine acquisition and mitigates sorafenib-induced ferroptosis in hepatocellular carcinoma. J Exp Clin Cancer Res 41, 98. 10.1186/s13046-022-02296-3.

Chen, M., Zhu, J.-Y., Mu, W.-J., and Guo, L. (2022). Cysteine dioxygenase type 1 (CDO1): Its functional role in physiological and pathophysiological processes. Genes & Diseases. https://doi.org/10.1016/j.gendis.2021.12.023.

Choi, J.I., Cho, E.H., Kim, S.B., Kim, R., Kwon, J., Park, M., Shin, H.J., Ryu, H.S., Park, S.H., and Lee, K.H. (2017). Promoter methylation of cysteine dioxygenase type 1: gene silencing and tumorigenesis in hepatocellular carcinoma. Ann Hepatobiliary Pancreat Surg 21, 181–187. 10.14701/ahbps.2017.21.4.181.

Combs, J.A., and DeNicola, G.M. (2019). The Non-Essential Amino Acid Cysteine Becomes Essential for Tumor Proliferation and Survival. Cancers 11. 10.3390/cancers11050678.

Commisso, C., Davidson, S.M., Soydaner-Azeloglu, R.G., Parker, S.J., Kamphorst, J.J., Hackett, S., Grabocka, E., Nofal, M., Drebin, J.A., Thompson, C.B., et al. (2013). Macropinocytosis of protein is an amino acid supply route in Ras-transformed cells. Nature 497, 633–637. 10.1038/nature12138.

Cramer, S.L., Saha, A., Liu, J., Tadi, S., Tiziani, S., Yan, W., Triplett, K., Lamb, C., Alters, S.E., and Rowlinson, S. (2017). Systemic depletion of L-cyst (e) ine with cyst (e) inase increases reactive oxygen species and suppresses tumor growth. Nature medicine 23, 120.

DeBlasi, J.M., Falzone, A., Caldwell, S., Kang, Y.P., Prieto-Farigua, N., Prigge, J.R., Schmidt, E.E., Chio, I.I.C., Karreth, F.A., and DeNicola, G.M. (2022). Distinct Nrf2 Signaling Thresholds Mediate Lung Tumor Initiation and Progression. bioRxiv, 2022.2008.2024.504986. 10.1101/2022.08.24.504986.

Dixon, S.J., Lemberg, K.M., Lamprecht, M.R., Skouta, R., Zaitsev, E.M., Gleason, C.E., Patel, D.N., Bauer, A.J., Cantley, A.M., Yang, W.S., et al. (2012). Ferroptosis: an iron-dependent form of nonapoptotic cell death. Cell 149, 1060–1072. 10.1016/j.cell.2012.03.042.

Gu, Y., Albuquerque, C.P., Braas, D., Zhang, W., Villa, G.R., Bi, J., Ikegami, S., Masui, K., Gini, B., Yang, H., et al. (2017). mTORC2 Regulates Amino Acid Metabolism in Cancer by Phosphorylation of the Cystine-Glutamate Antiporter xCT. Molecular Cell 67, 128–138.e127. https://doi.org/10.1016/j.molcel.2017.05.030.

Harris, I.S., and DeNicola, G.M. (2020). The Complex Interplay between Antioxidants and ROS in Cancer. Trends in Cell Biology.

Helms, E., Onate, M.K., and Sherman, M.H. (2020). Fibroblast Heterogeneity in the Pancreatic Tumor Microenvironment. Cancer Discov 10, 648–656. 10.1158/2159-8290.Cd-19-1353.

Igarashi, K., Yamashita, K., Katoh, H., Kojima, K., Ooizumi, Y., Nishizawa, N., Nishiyama, R., Kawamata, H., Tajima, H., Kaizu, T., et al. (2017). Prognostic significance of promoter DNA hypermethylation of the cysteine dioxygenase 1 (CDO1) gene in primary gallbladder cancer and gallbladder disease. PLoS One 12, e0188178. 10.1371/journal.pone.0188178.

Jackson, E.L., Olive, K.P., Tuveson, D.A., Bronson, R., Crowley, D., Brown, M., and Jacks, T. (2005). The differential effects of mutant p53 alleles on advanced murine lung cancer. Cancer Res 65, 10280–10288. 10.1158/0008-5472.Can-05-2193.

Jiang, L., Kon, N., Li, T., Wang, S.J., Su, T., Hibshoosh, H., Baer, R., and Gu, W. (2015). Ferroptosis as a p53-mediated activity during tumour suppression. Nature 520, 57–62. 10.1038/nature14344.

Kang, Y.P., Mockabee-Macias, A., Jiang, C., Falzone, A., Prieto-Farigua, N., Stone, E., Harris, I.S., and DeNicola, G.M. (2021). Non-canonical Glutamate-Cysteine Ligase Activity Protects against Ferroptosis. Cell Metab 33, 174–189.e177. 10.1016/j.cmet.2020.12.007.

Kang, Y.P., Torrente, L., Falzone, A., Elkins, C.M., Liu, M., Asara, J.M., Dibble, C.C., and DeNicola, G.M. (2019). Cysteine dioxygenase 1 is a metabolic liability for non-small cell lung cancer. eLife 8, e45572. 10.7554/eLife.45572.

Kim, J., Hong, S.J., Park, J.H., Park, S.Y., Kim, S.W., Cho, E.Y., Do, I.G., Joh, J.W., and Kim, D.S. (2009). Expression of cystathionine beta-synthase is downregulated in hepatocellular carcinoma and associated with poor prognosis. Oncol Rep 21, 1449–1454. 10.3892/or_00000373.

Kojima, K., Nakamura, T., Ohbu, M., Katoh, H., Ooizumi, Y., Igarashi, K., Ishii, S., Tanaka, T., Yokoi, K., Nishizawa, N., et al. (2018). Cysteine dioxygenase type 1 (CDO1) gene promoter methylation during the adenoma-carcinoma sequence in colorectal cancer. PLoS One 13, e0194785. 10.1371/journal.pone.0194785.

Lien, E.C., Ghisolfi, L., Geck, R.C., Asara, J.M., and Toker, A. (2017). Oncogenic PI3K promotes methionine dependency in breast cancer cells through the cystine-glutamate antiporter xCT. Sci Signal 10. 10.1126/scisignal.aao6604.

Lin, Z., Huang, W., He, Q., Li, D., Wang, Z., Feng, Y., Liu, D., Zhang, T., Wang, Y., Xie, M., et al. (2021). FOXC1 promotes HCC proliferation and metastasis by Upregulating DNMT3B to induce DNA Hypermethylation of CTH promoter. J Exp Clin Cancer Res 40, 50. 10.1186/s13046-021-01829-6.

Lu, S.C. (1999). Regulation of hepatic glutathione synthesis: current concepts and controversies. Faseb j 13, 1169–1183.

Maekawa, H., Ito, T., Orita, H., Kushida, T., Sakurada, M., Sato, K., Hulbert, A., and Brock, M.V. (2020). Analysis of the methylation of CpG islands in the CDO1, TAC1 and CHFR genes in pancreatic ductal cancer. Oncol Lett 19, 2197–2204. 10.3892/ol.2020.11340.

Masson, N., Keeley, T.P., Giuntoli, B., White, M.D., Puerta, M.L., Perata, P., Hopkinson, R.J., Flashman, E., Licausi, F., and Ratcliffe, P.J. (2019). Conserved N-terminal cysteine dioxygenases transduce responses to hypoxia in animals and plants. Science 365, 65–69. 10.1126/science.aaw0112.

Mollaoglu, G., Guthrie, M.R., Böhm, S., Brägelmann, J., Can, I., Ballieu, P.M., Marx, A., George, J., Heinen, C., Chalishazar, M.D., et al. (2017). MYC Drives Progression of Small Cell Lung Cancer to a Variant Neuroendocrine Subtype with Vulnerability to Aurora Kinase Inhibition. Cancer Cell 31, 270–285. 10.1016/j.ccell.2016.12.005.

Morton, J.P., Timpson, P., Karim, S.A., Ridgway, R.A., Athineos, D., Doyle, B., Jamieson, N.B., Oien, K.A., Lowy, A.M., Brunton, V.G., et al. (2010). Mutant p53 drives metastasis and overcomes growth arrest/senescence in pancreatic cancer. Proc Natl Acad Sci U S A 107, 246–251. 10.1073/pnas.0908428107.

Mudd, S.H., Finkelstein, J.D., Irreverre, F., and Laster, L. (1965). Transsulfuration in mammals. Microassays and tissue distributions of three enzymes of the pathway. J Biol Chem 240, 4382–4392.

Mukhopadhyay, S., Biancur, D.E., Parker, S.J., Yamamoto, K., Banh, R.S., Paulo, J.A., Mancias, J.D., and Kimmelman, A.C. (2021). Autophagy is required for proper cysteine homeostasis in pancreatic cancer through regulation of SLC7A11. Proc Natl Acad Sci U S A 118. 10.1073/pnas.2021475118.

Nishimura, T., Higuchi, K., Yoshida, Y., Sugita-Fujisawa, Y., Kojima, K., Sugimoto, M., Santo, M., Tomi, M., and Nakashima, E. (2018). Hypotaurine Is a Substrate of GABA Transporter Family Members GAT2/Slc6a13 and TAUT/Slc6a6. Biol Pharm Bull 41, 1523–1529. 10.1248/bpb.b18-00168.

Nishizawa, N., Harada, H., Kumamoto, Y., Kaizu, T., Katoh, H., Tajima, H., Ushiku, H., Yokoi, K., Igarashi, K., Fujiyama, Y., et al. (2019). Diagnostic potential of hypermethylation of the cysteine dioxygenase 1 gene (CDO1) promoter DNA in pancreatic cancer. Cancer Sci 110, 2846–2855. 10.1111/cas.14134.

Ookhtens, M., and Kaplowitz, N. (1998). Role of the Liver in Interorgan Homeostasis of Glutathione and Cyst(e)ine. Semin Liver Dis 18, 313–329.

Ooki, A., Maleki, Z., Tsay, J.J., Goparaju, C., Brait, M., Turaga, N., Nam, H.S., Rom, W.N., Pass, H.I., Sidransky, D., et al. (2017). A Panel of Novel Detection and Prognostic Methylated DNA Markers in Primary Non-Small Cell Lung Cancer and Serum DNA. Clin Cancer Res 23, 7141–7152. 10.1158/1078-0432.Ccr-17-1222.

Orsatti, L., Orsale, M.V., di Pasquale, P., Vecchi, A., Colaceci, F., Ciammaichella, A., Rossetti, I., Bonelli, F., Baumgaertel, K., Liu, K., et al. (2021). Turnover rate of coenzyme A in mouse brain and liver. PLoS One 16, e0251981. 10.1371/journal.pone.0251981.

Pavlova, N.N., and Thompson, C.B. (2016). The Emerging Hallmarks of Cancer Metabolism. Cell Metab 23, 27–47. 10.1016/j.cmet.2015.12.006.

Pavlova, N.N., Zhu, J., and Thompson, C.B. (2022). The hallmarks of cancer metabolism: Still emerging. Cell Metab 34, 355–377. 10.1016/j.cmet.2022.01.007.

Ruiz de Galarreta, M., Bresnahan, E., Molina-Sánchez, P., Lindblad, K.E., Maier, B., Sia, D., Puigvehi, M., Miguela, V., Casanova-Acebes, M., Dhainaut, M., et al. (2019). β-Catenin Activation Promotes Immune Escape and Resistance to Anti-PD-1 Therapy in Hepatocellular Carcinoma. Cancer Discov 9, 1124–1141. 10.1158/2159-8290.Cd-19-0074.

Sasaki, H., Sato, H., Kuriyama-Matsumura, K., Sato, K., Maebara, K., Wang, H., Tamba, M., Itoh, K., Yamamoto, M., and Bannai, S. (2002). Electrophile response element-mediated induction of the cystine/glutamate exchange transporter gene expression. J Biol Chem 277, 44765–44771. 10.1074/jbc.M208704200.

Sato, H., Nomura, S., Maebara, K., Sato, K., Tamba, M., and Bannai, S. (2004). Transcriptional control of cystine/glutamate transporter gene by amino acid deprivation. Biochem Biophys Res Commun 325, 109–116. 10.1016/j.bbrc.2004.10.009.

Sato, H., Shiiya, A., Kimata, M., Maebara, K., Tamba, M., Sakakura, Y., Makino, N., Sugiyama, F., Yagami, K., Moriguchi, T., et al. (2005). Redox imbalance in cystine/glutamate transporter-deficient mice. J Biol Chem 280, 37423–37429. 10.1074/jbc.M506439200.

Stipanuk, M.H. (2004a). Role of the liver in regulation of body cysteine and taurine levels: a brief review. Neurochem Res 29, 105–110. 10.1023/b:nere.0000010438.40376.c9.

Stipanuk, M.H. (2004b). Sulfur amino acid metabolism: pathways for production and removal of homocysteine and cysteine. Annu Rev Nutr 24, 539–577. 10.1146/annurev.nutr.24.012003.132418.

Sullivan, M.R., Danai, L.V., Lewis, C.A., Chan, S.H., Gui, D.Y., Kunchok, T., Dennstedt, E.A., Vander Heiden, M.G., and Muir, A. (2019). Quantification of microenvironmental metabolites in murine cancers reveals determinants of tumor nutrient availability. Elife 8. 10.7554/eLife.44235.

Sumizu, K. (1962). Oxidation of hypotaurine in rat liver. Biochim Biophys Acta 63, 210–212. 10.1016/0006-3002(62)90357-8.

Tsuchihashi, K., Okazaki, S., Ohmura, M., Ishikawa, M., Sampetrean, O., Onishi, N., Wakimoto, H., Yoshikawa, M., Seishima, R., Iwasaki, Y., et al. (2016). The EGF Receptor Promotes the Malignant Potential of Glioma by Regulating Amino Acid Transport System xc(-). Cancer Res 76, 2954–2963. 10.1158/0008-5472.Can-15-2121.

Veeravalli, S., Phillips, I.R., Freire, R.T., Varshavi, D., Everett, J.R., and Shephard, E.A. (2020). Flavin-Containing Monooxygenase 1 Catalyzes the Production of Taurine from Hypotaurine. Drug Metab Dispos 48, 378–385. 10.1124/dmd.119.089995.

Wang, L., Han, H., Liu, Y., Zhang, X., Shi, X., and Wang, T. (2018). Cystathionine β-synthase Induces Multidrug Resistance and Metastasis in Hepatocellular Carcinoma. Curr Mol Med 18, 496–506. 10.2174/1566524019666181211162754.

Wang, L., Xing, X., Zeng, X., Jackson, S.R., TeSlaa, T., Al-Dalahmah, O., Samarah, L.Z., Goodwin, K., Yang, L., McReynolds, M.R., et al. (2022). Spatially resolved isotope tracing reveals tissue metabolic activity. Nat Methods 19, 223–230. 10.1038/s41592-021-01378-y.

Ward, N.P., and DeNicola, G.M. (2019). Chapter Three - Sulfur metabolism and its contribution to malignancy. In International Review of Cell and Molecular Biology, D.C. Montrose, and L. Galluzzi, eds. (Academic Press), pp. 39–103.https://doi.org/10.1016/bs.ircmb.2019.05.001.

Winterbourn, C.C., and Hampton, M.B. (2008). Thiol chemistry and specificity in redox signaling. Free Radical Biology and Medicine 45, 549–561.

Ye, C., Sutter, B.M., Wang, Y., Kuang, Z., and Tu, B.P. (2017). A Metabolic Function for Phospholipid and Histone Methylation. Mol Cell 66, 180–193.e188. 10.1016/j.molcel.2017.02.026.

Yin, W., Wang, X., Li, Y., Wang, B., Song, M., Hulbert, A., Chen, C., and Yu, F. (2020). Promoter hypermethylation of cysteine dioxygenase type 1 in patients with non-small cell lung cancer. Oncol Lett 20, 967–973. 10.3892/ol.2020.11592.

Zhang, H.F., Hughes, C.S., Li, W., He, J.Z., Surdez, D., El-Naggar, A.M., Cheng, H., Prudova, A., Delaidelli, A., Negri, G.L., et al. (2021). Proteomic Screens for Suppressors of Anoikis Identify IL1RAP as a Promising Surface Target in Ewing Sarcoma. Cancer Discov 11, 2884–2903. 10.1158/2159-8290.Cd-20-1690.

Zhang, H.F., Klein Geltink, R.I., Parker, S.J., and Sorensen, P.H. (2022). Transsulfuration, minor player or crucial for cysteine homeostasis in cancer. Trends Cell Biol. 10.1016/j.tcb.2022.02.009.

Zhang, T., Bauer, C., Newman, A.C., Uribe, A.H., Athineos, D., Blyth, K., and Maddocks, O.D.K. (2020). Polyamine pathway activity promotes cysteine essentiality in cancer cells. Nat Metab 2, 1062–1076. 10.1038/s42255-020-0253-2.

Zhang, Y., Tan, H., Daniels, J.D., Zandkarimi, F., Liu, H., Brown, L.M., Uchida, K., O’Connor, O.A., and Stockwell, B.R. (2019). Imidazole ketone erastin induces ferroptosis and slows tumor growth in a mouse lymphoma model. Cell chemical biology 26, 623–633. e629.

Zhou, Y., and Danbolt, N.C. (2013). GABA and Glutamate Transporters in Brain. Front Endocrinol (Lausanne) 4, 165. 10.3389/fendo.2013.00165.

Zhu, J., Berisa, M., Schwörer, S., Qin, W., Cross, J.R., and Thompson, C.B. (2019). Transsulfuration Activity Can Support Cell Growth upon Extracellular Cysteine Limitation. Cell Metab 30, 865–876.e865. 10.1016/j.cmet.2019.09.009.

