## Supplemental Information for "The Origin of Cysteine and its Catabolism in Mammalian Tissues and Tumors"

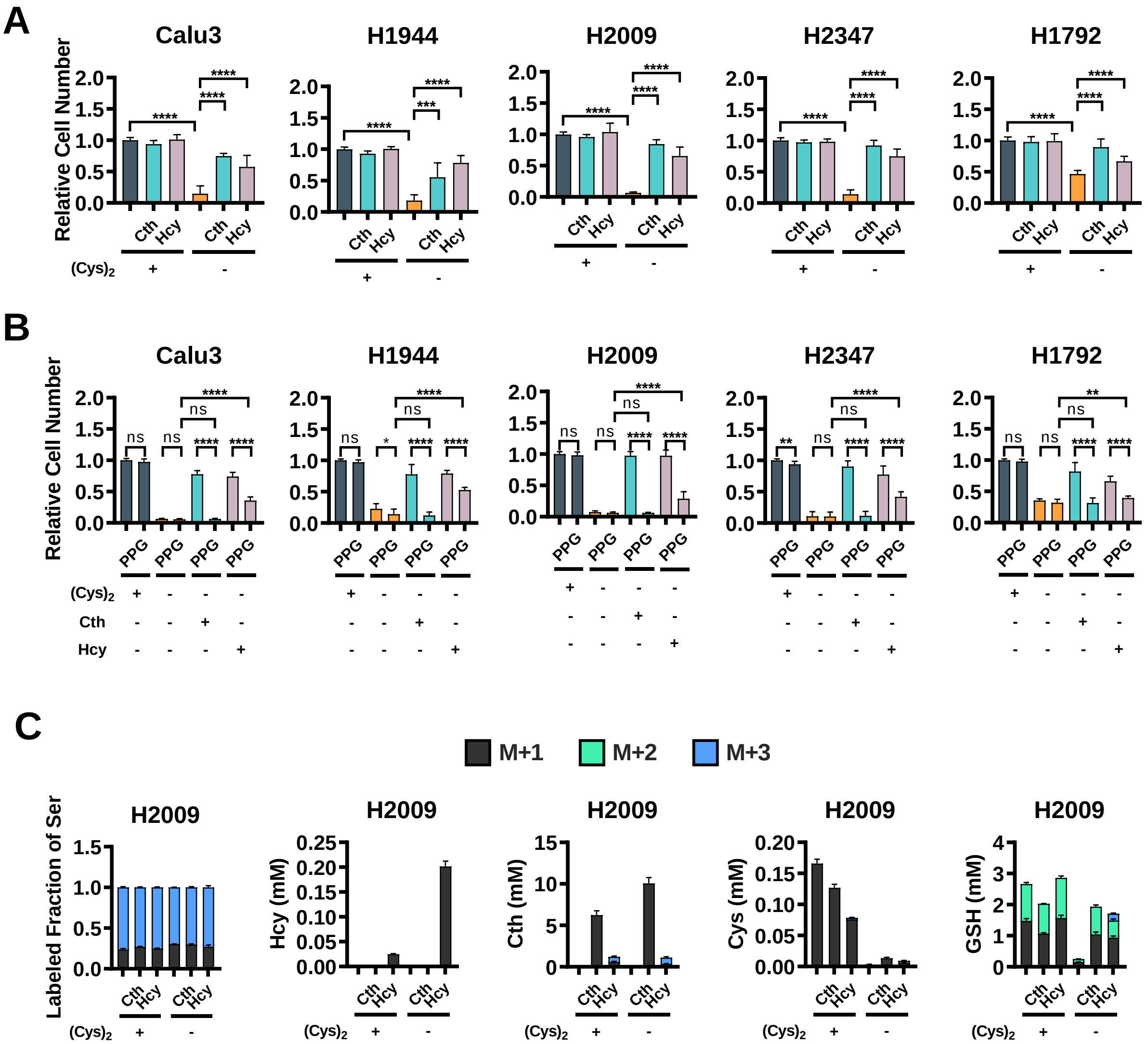
A
Calu3
H1944
H2009
H2347
H1792
Relative Cell Number
Cth
Hcy
(Cys)2
+
-
B
Calu3
H1944
H2009
H2347
H1792
Relative Cell Number
PPG
ns
(Cys)2
Cth
Hcy
C
M+1
M+2
M+3
H2009
Labeled Fraction of Ser
Hcy (mM)
Cth (mM)
Cys (mM)
GSH (mM)

**Figure S1. Cystathionine and Homocysteine Rescue NSCLC Cell Viability under Cystine Starvation (related to Figure 1).**

(A) Measurement of relative NSCLC cell number under cystine-replete (200 μM) or cystine-starved (0 μM) conditions supplemented with 1 mM cystathionine or 1 mM homocysteine. (B) Measurement of relative NSCLC cell number under cystine-starved condition treated with cystathionine or homocysteine in the presence of absence of 100 μM of propargylglycine. Relative cell number was determined by crystal violet staining, followed by normalization to cystine-replete, untreated conditions. For (A) and (B), data are presented as mean ± SD. (C) Labeled fraction of serine and concentration of homocysteine, cystathionine, cysteine, and glutathione after 24 h $^{13}C_3$-serine labeling in H2009 under cystine-replete (200 μM) or cystine-starved (0 μM) condition supplemented with 1 mM cystathionine or 1 mM homocysteine. N=3 biological replicates for each condition. *$p < 0.05$, **$p < 0.01$, ***$p < 0.001$, and ****$p < 0.0001$. Cys, cysteine; $(Cys)_2$, cystine; Cth, cystathionine; Hcy, homocysteine; PPG, propargylglycine; GSH, glutathione; Ser, serine.

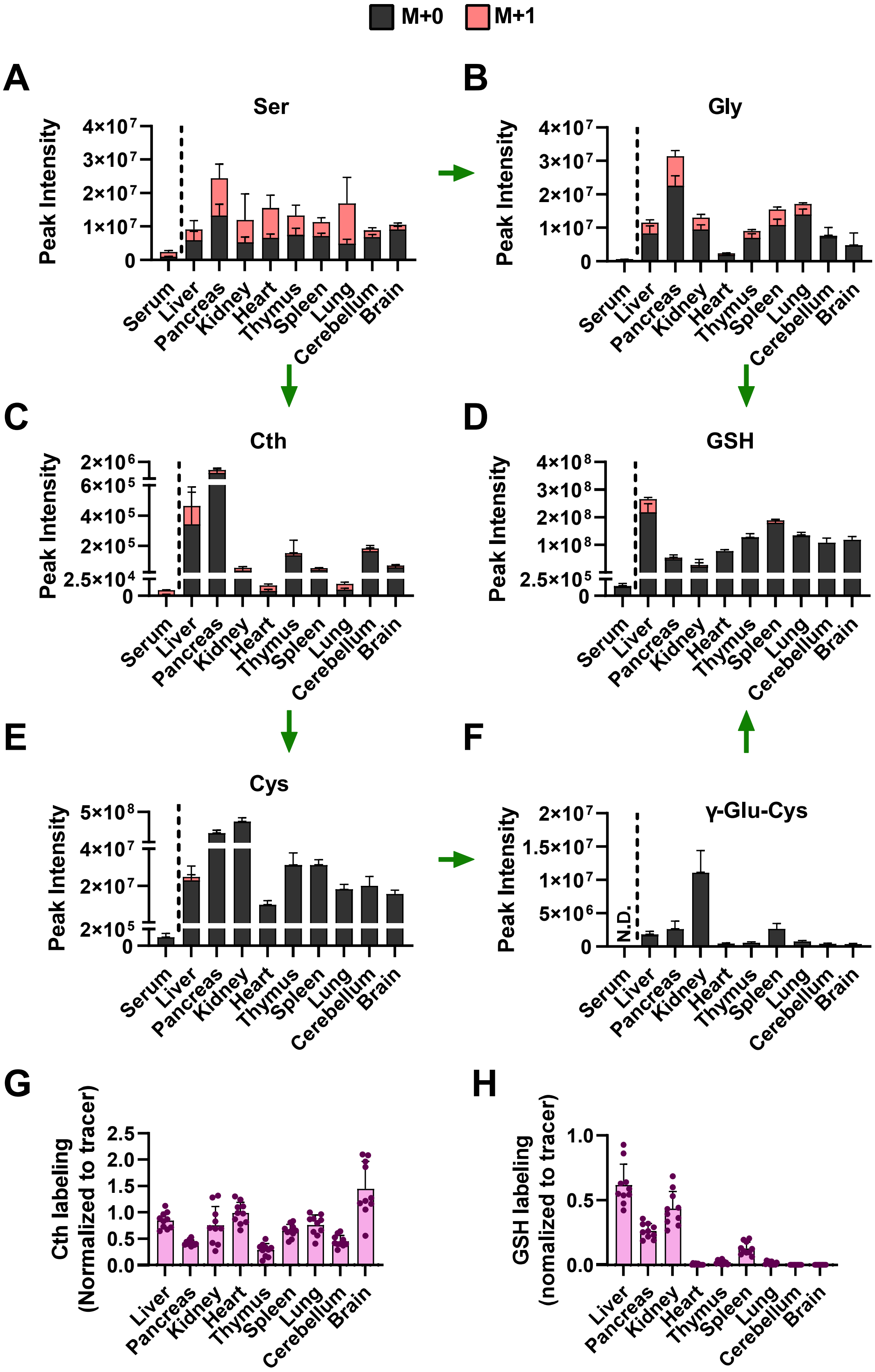

M+0
M+1
A
Ser
Peak Intensity
4×10^7
3×10^7
2×10^7
1×10^7
0
Serum
Liver
Pancreas
Kidney
Heart
Thymus
Spleen
Lung
Cerebellum
Brain
B
Gly
Peak Intensity
4×10^7
3×10^7
2×10^7
1×10^7
0
C
Cth
Peak Intensity
2×10^6
6×10^5
4×10^5
2×10^5
2.5×10^4
0
D
GSH
4×10^8
3×10^8
2×10^8
1×10^8
2.5×10^5
0
E
Cys
Peak Intensity
5×10^8
4×10^7
2×10^7
2×10^5
0
F
γ-Glu-Cys
Peak Intensity
2×10^7
1.5×10^7
1×10^7
5×10^6
0
N.D.
G
Cth labeling
(Normalized to tracer)
2.5
2.0
1.5
1.0
0.5
0.0
H
GSH labeling
(nomalized to tracer)
1.0
0.5
0.0

**Figure S2. Total levels of serine, cysteine and glutathione intermediates in healthy mouse tissues (related to Figure 2).**

(A-F) Healthy C57BL6J mice were infused with 1-[$^{13}C_3$]-serine for 4 hours, followed by analysis of the total signal of (A) serine, (B) glycine, (C) cystathionine, (D) glutathione, (E) cysteine and (F) γ-glutamylcysteine in the serum and nine different tissues (liver, pancreas, kidney, heart, thymus, spleen, lung, cerebellum, and brain) (N=10 mice). The total signal is the sum of all isotopologue signals. The peak intensities of isotopologues were measured as AreaTop (mean of three top points in the peak). (G-H) Fractional contribution of serine to (G) cystathionine and (H) glutathione in the tissues from (A-F). Metabolite labeling was normalized to the fraction labeling of serine in each tissue. For (A-H), data are presented as mean ± SD. N.D., not detected. Ser, serine; Cth, cystathionine; Cys, cysteine; Gly, glycine; GSH, glutathione; γ-Glu-Cys, γ-glutamylcysteine.

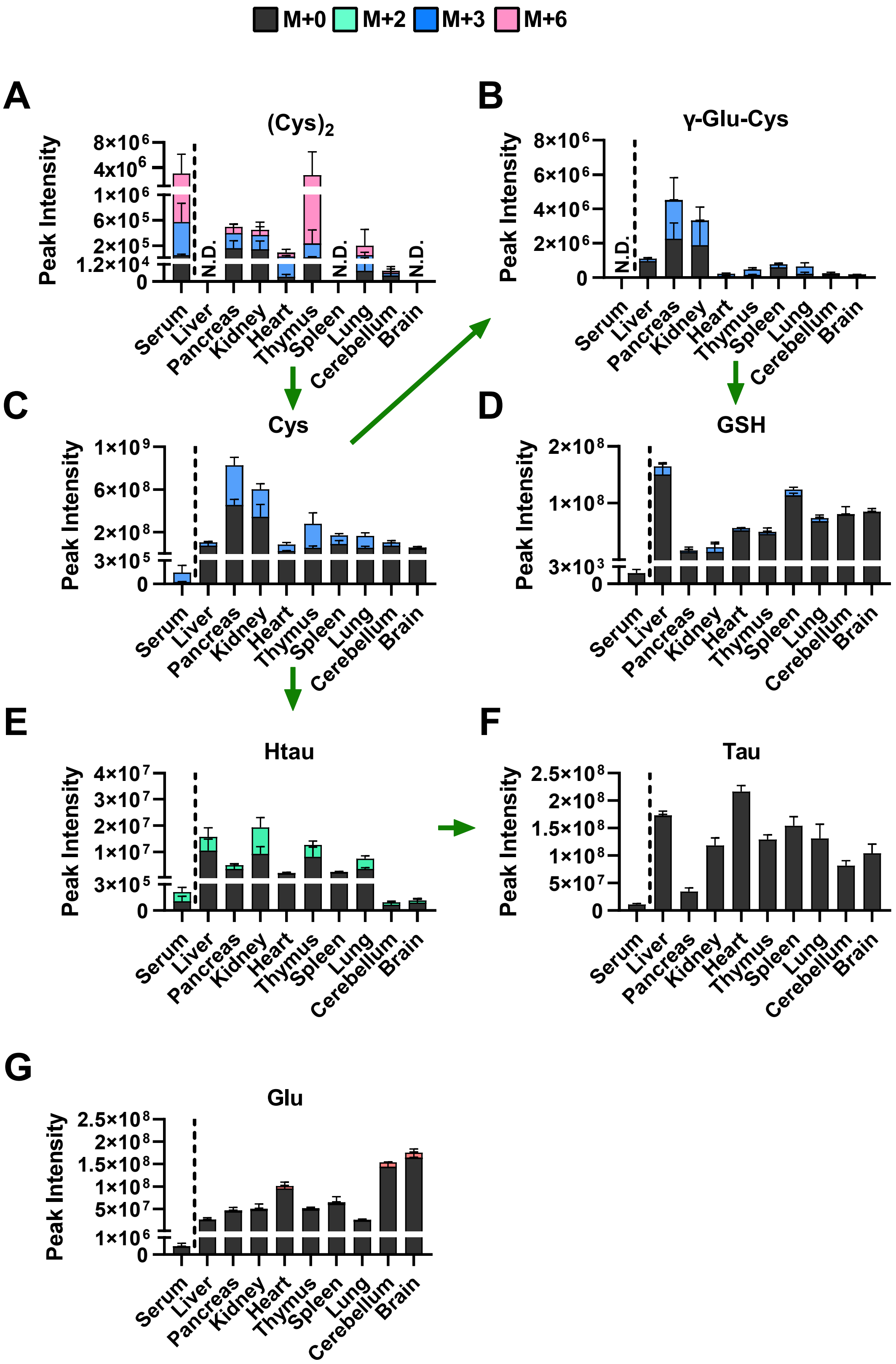
M+0
M+2
M+3
M+6
A
$(Cys)_2$
B
γ-Glu-Cys
C
Cys
D
GSH
E
Htau
F
Tau
G
Glu
Peak Intensity
N.D.
Serum
Liver
Pancreas
Kidney
Heart
Thymus
Spleen
Lung
Cerebellum
Brain

**Figure S3. Total levels of cysteine, glutathione, and taurine intermediates in healthy mouse tissues (related to Figure 3).**

(A-F) Healthy C57BL6J mice were infused with $^{13}C_6$-cystine for 4 hours, followed by analysis of the total signal of (A) cystine, (B) γ-glutamylcysteine, (C) cysteine, (D) glutathione, (E) hypotaurine and (F) taurine. For (A-F), data are presented as mean ± SD and N=5 mice. The total signal is the sum of all isotopologue signals. The peak intensities of isotopologues were measured as AreaTop (mean of three top points in the peak). (G) Fractional contribution of cystine to γ-glutamylcysteine in each tissue from (A-F). γ-glutamylcysteine labeling was normalized to the fraction labeling of cysteine in each tissue. Ser, serine; Cth, cystathionine; Cys, cysteine; Gly, glycine; GSH, glutathione; γ-Glu-Cys, γ-glutamylcysteine. N.D., not detected.

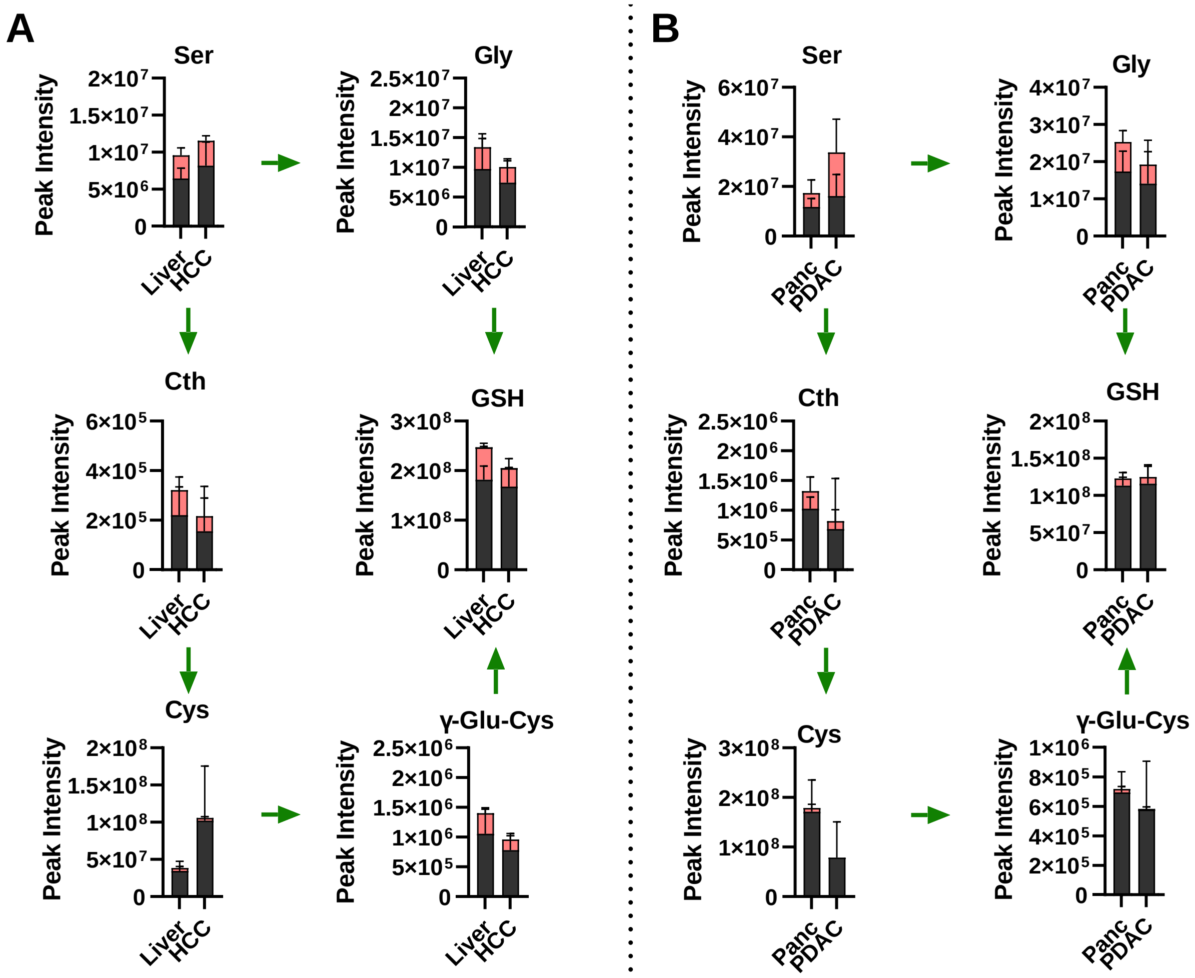
A
Ser
Gly
Cth
GSH
Cys
γ-Glu-Cys
Peak Intensity
Liver
HCC
B
Panc
PDAC

**Figure S4. Total levels of cysteine and glutathione intermediates in liver and pancreas cancer models (related to Figure 4).**

(A) Analysis of the total signal of serine, glycine, cystathionine, glutathione, cysteine and γ-glutamylcysteine in control liver (N=8) and HCC (N=8) following infusion with 1-[$^{13}C_3$]-serine. (B) Analysis of the total signal of = serine, glycine, cystathionine, glutathione, cysteine and γ-glutamylcysteine in in control pancreas (N=5) and PDAC (N=5) following infusion with 1-[$^{13}C_3$]-serine. N.B. the control lung samples in (A) are the same as in (B). Data are presented as mean ± SD. The total signal is the sum of all isotopologue signals. The peak intensities of isotopologues were measured as AreaTop (mean of three top points in the peak). Ser, serine; Cth, cystathionine; Cys, cysteine; Gly, glycine; GSH, glutathione; γ-Glu-Cys, γ-glutamylcysteine

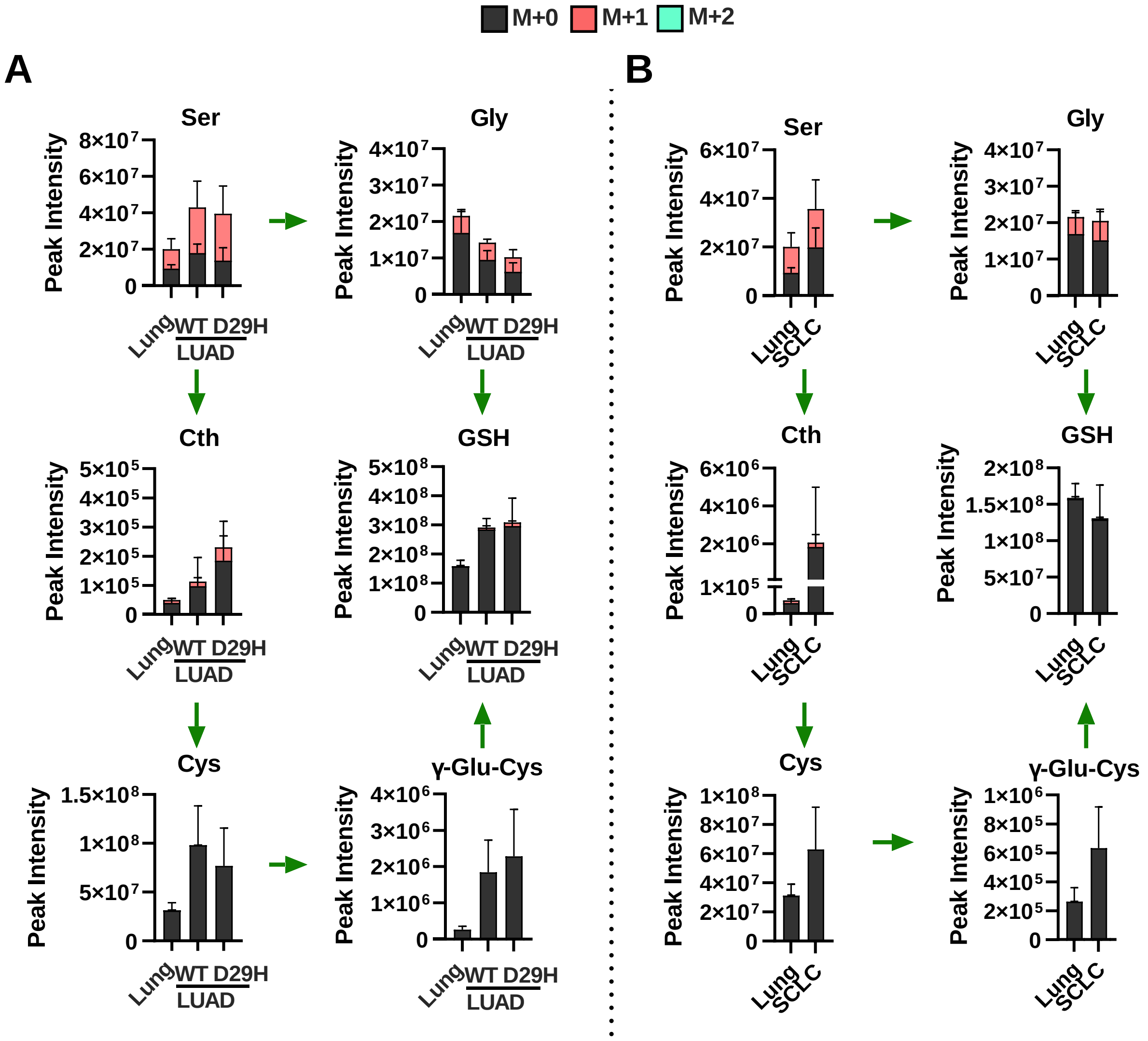

M+0
M+1
M+2
A
B
Ser
Gly
Cth
GSH
Cys
γ-Glu-Cys
Peak Intensity
Lung
WT
D29H
LUAD
SCLC

**Figure S5. Total levels of cysteine and glutathione intermediates in lung cancer models (related to Figure 5).**

(A) Analysis of the total signal of serine, glycine, cystathionine, glutathione, cysteine and γ-glutamylcysteine in control lung (N=8) and Nrf2$^{WT}$ lung adenocarcinoma (N=10) and Nrf2$^{D29H}$ lung adenocarcinoma (N=10) following infusion with 1-[$^{13}C_3$]-serine. (B) Analysis of the total signal of = serine, glycine, cystathionine, glutathione, cysteine and γ-glutamylcysteine in in control lung (N=8) and small cell lung cancer (SCLC, N=10) following infusion with 1-[$^{13}C_3$]-serine. N.B. the control lung samples in (A) are the same as in (B). Data are presented as mean ± SD. The total signal is the sum of all isotopologue signals. The peak intensities of isotopologues were measured as AreaTop (mean of three top points in the peak). Ser, serine; Cth, cystathionine; Cys, cysteine; Gly, glycine; GSH, glutathione; γ-Glu-Cys, γ-glutamylcysteine

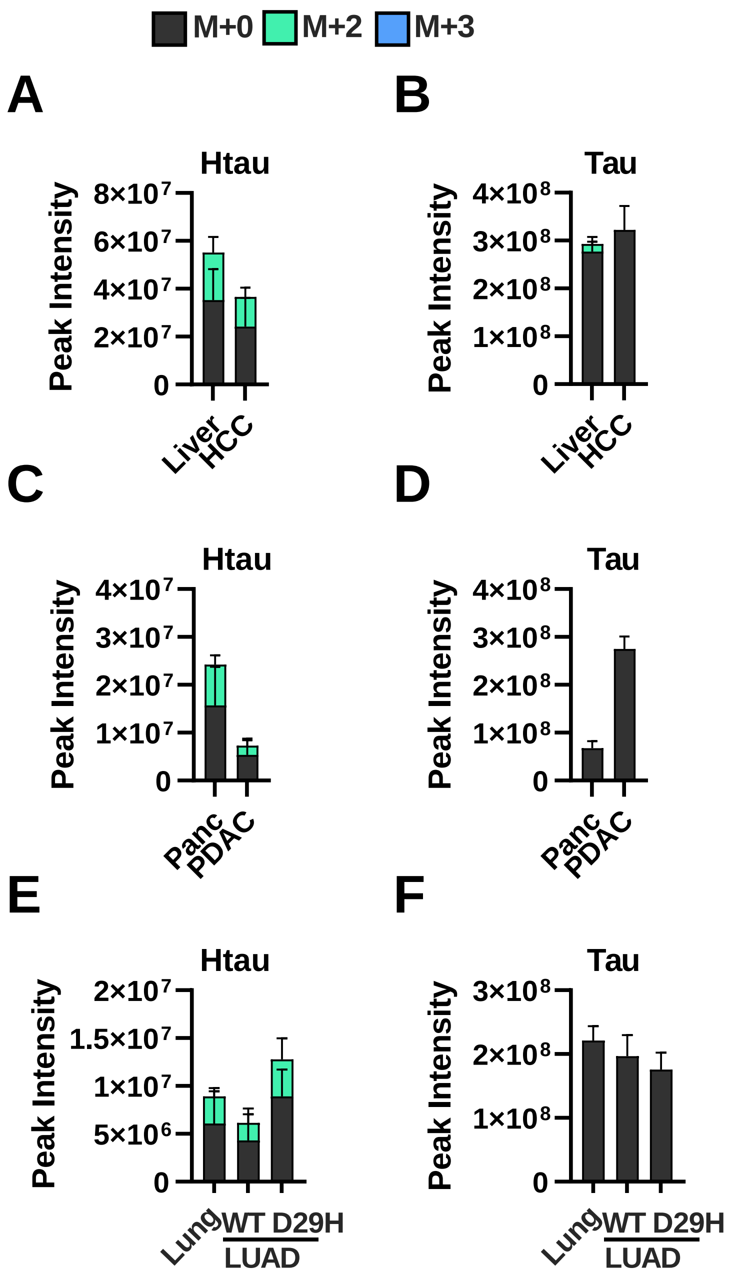

M+0
M+2
M+3
A
Htau
Peak Intensity
8×10⁷
6×10⁷
4×10⁷
2×10⁷
0
Liver
HCC
B
Tau
Peak Intensity
4×10⁸
3×10⁸
2×10⁸
1×10⁸
0
Liver
HCC
C
Htau
Peak Intensity
4×10⁷
3×10⁷
2×10⁷
1×10⁷
0
Panc
PDAC
D
Tau
Peak Intensity
4×10⁸
3×10⁸
2×10⁸
1×10⁸
0
Panc
PDAC
E
Htau
Peak Intensity
2×10⁷
1.5×10⁷
1×10⁷
5×10⁶
0
Lung
WT
D29H
LUAD
F
Tau
Peak Intensity
3×10⁸
2×10⁸
1×10⁸
0
Lung
WT
D29H
LUAD

**Figure S6. Total levels of taurine pathway intermediates in liver, pancreas, and lung cancer models (related to Figure 7).**

(A-F) Analysis of the total signal of hypotaurine (A, C, E) and taurine (B, D, E) in control liver (N=6) and HCC (N=4) (A, B), control pancreas (N=3) and PDAC (N=12) (C, D) and control lung (N=10), $Nrf2^{WT}$ LUAD (N=16), and $Nrf2^{D29H}$ LUAD (N=10) (E, F) following infusion with $^{13}C_6$-cystine. Data are presented as mean ± SD. The total signal is the sum of all isotopologue signals. The peak intensities of isotopologues were measured as AreaTop (mean of three top points in the peak). Htau, hypotaurine; Tau, taurine
